# IFNγ and proinflammatory cytokines cooperatively license astrocytes to support antigen-dependent CD4+ T cell restimulation

**DOI:** 10.64898/2026.07.29.741554

**Authors:** Theodore M Fisher, Francesco Limone, Matthew D Smith, Chandler Wright, Petra Kukanja, Peiwen Lu, Kerry Limberg, Nicolette Pirjanian, Jimena Zulueta, Valentina Fossati, Akiko Iwasaki, Peter A Calabresi, Shane A Liddelow

## Abstract

We report that type 2 interferon with proinflammatory cytokines induces astrocytes to express multiple components required for attraction of and antigen presentation to CD4+ T cells via MHC-II molecules. Re-analysis of astrocyte-enriched single-cell sequencing from bacterial mimetic-injected mice revealed a transcriptionally distinct interferon-responsive astrocyte population enriched for antigen-presentation-associated genes 24 hours after peripheral immune challenge. In vivo, we identify MHC-II+GFAP+ astrocytes localized to the surface of the brain across multiple mouse models of neuroinflammation and in human neurological disease tissue. In a pure in vitro culture system, astrocytes stimulated with IFNγ and proinflammatory cytokines express MHC-II, costimulatory molecules and lymphocyte-recruiting chemokines, and support rapid antigen-dependent expansion of previously activated CD4+ T cells. Using human iPSC-derived CNS cultures, we additionally show induction of antigen-presentation-associated machinery in human astrocytes following inflammatory cytokine stimulation. Rather than inferring function solely from marker expression, we assessed astrocyte-supported T cell restimulation in vitro and separately characterized the emergence of antigen-presentation-associated astrocytes in vivo at neuroimmune interfaces during inflammation in mice and humans. Together, these findings support a conserved role for cytokine-stimulated astrocytes in expressing antigen-presentation-associated machinery and, in vitro, supporting antigen-dependent restimulation/expansion of CD4+ T cells under inflammatory conditions.

## INTRODUCTION

Astrocytes, long appreciated for their supportive roles in CNS homeostasis, are increasingly recognized as dynamic immune effectors during neuroinflammation^1-3^. Under pathological conditions, astrocytes become reactive, acquiring transcriptionally heterogeneous identities and diverse phenotypes that influence various functions in the CNS^4,5^. Among their many emerging roles, recent evidence highlights the capacity of reactive astrocytes to interface with and modulate lymphocytes through cytokine release^6,7^ and antigen presentation^8-13^, directly influencing CNS coordination of adaptive immunity. Although prior studies have reported expression of MHC-II-related molecules by astrocytes, the cytokine requirements for induction of a broader antigen-presentation program in astrocytes, and the capacity of this program to support antigen-specific CD4+ T cell responses, remain incompletely defined.

While the implication of reactive astrocytes in chemokine and cytokine axes have been investigated extensively, important questions remain regarding the cytokine requirements that induce a broader antigen-presentation program in astrocytes and the extent to which stimulated astrocytes can functionally support antigen-specific CD4+ T cell responses. Prior studies have reported increased MHC-II-related transcription and protein levels by astrocytes across multiple inflammatory and neurodegenerative contexts^4,14-17^, including after IFNγ exposure, but the signaling logic required to induce a more complete antigen-presentation phenotype and its functional consequences for T cell restimulation remain incompletely defined.

Astrocyte-specific type II major histocompatibility complex (MHC-II)-related gene transcription has been reported in multiple studies in the contexts of Alzheimer’s disease (AD)^14,15^, infection^4,16^, CNS injury^17^, multiple sclerosis (MS)^18-20^, and Parkinson’s disease^10^. MHC-II protein production by astrocytes has also been shown in vitro after treatment with IFNγ^10,12^. However, the presence of MHC-II is only one piece of the complex antigen-presenting machinery required for appropriate processing and presentation of antigens to immune cells. In this study, we address a key gap in understanding the molecular stimuli required to induce MHC-II and related machinery needed for functional interaction with CD4+ T cells.

Antigen presentation is a central immune communication process conserved in mammals that influences adaptive immunity^21^. Presentation of peptide antigens by MHC molecules to T lymphocytes allows for the recognition of foreign and self-antigens. MHC class I displays peptides from intra-cellular proteins on the cell surface to communicate infection or damage to CD8+ T cells resulting in elimination^22^. Alternatively, MHC-II presents exogenous antigens sampled from the extra-cellular space to CD4+ helper T cells, a means of surveying and communicating immune challenges from the cellular environment^23^. This process requires coordinated antigen uptake, endosomal processing, and peptide loading onto MHC-II within the endolysosomal compartment, resulting in the membrane externalization of MHC complex and co-stimulatory molecules^24^. This mechanism is essential for initiating and shaping CD4+ T helper cell responses as circulating effector T cells rely on the antigen experience and inflammatory context delivered by antigen-presenting cells to promote tolerance, propagate inflammation, or generate memory populations during immune challenge^25^. While professional antigen presenters such as dendritic cells, macrophages, and B cells are classically responsible for MHC-II antigen presentation, accumulating evidence suggests that this process is more widely distributed, inducible, and dynamically regulated than previously appreciated^26^.

In atypical antigen presentation, mast cells, basophils, eosinophils, and innate lymphoid cells may acquire the machinery to present antigens under inflammatory or pathological conditions^27,28^. In these settings, presentation on MHC-II may be induced by cytokines such as interferon-gamma (IFNγ), allowing these cell types to directly interact with CD4+ T cells. Typically, initial recognition and relay of pathogenic insult to naïve T cells is performed by professional antigen presenting cells in the secondary lymphoid tissues including the spleen and lymph nodes. As these naïve T cells are stimulated and take on effector phenotypes, they enter circulation. Likely, atypical antigen presentation across various tissues provides a context-specific mechanism of providing antigen experience combinatorially with cytokine stimulation to instruct localized T cell responses^29,30^. Given its functional relevance to inflammatory propagation, atypical antigen presentation is increasingly recognized as a key feature of chronic inflammation, autoimmunity, and tissue-specific immune responses^27,31^. Consequently, considerable interest has emerged in determining whether other CNS-resident cell types –including microglia^32-34^, border macrophages^35,36^, and oligodendrocyte progenitor cells^37^ can similarly function as atypical antigen-presenting cells. Nevertheless, further functional studies are required to establish the extent and significance of antigen presentation across these CNS populations.

The CNS had long been considered immune-privileged until a paradigm shift with the discovery of immune processes performed by multiple glial and infiltrating immune cell populations in various neuroinflammatory contexts^1,2,4,38-40^. Concurrently, in many of these conditions, CD4+ T cells extravasate into the CNS and may expand clonally and locally^39,41-45^, but it remains unclear how they are able to do so. Still, it is unknown if they expand peripherally and migrate on mass into the CNS, or if they respond locally in the context pathologies like the demyelination that occurs in MS, around amyloid deposits in AD, or in response to degenerating neurons in amyotrophic lateral sclerosis.

Astrocytes are widely distributed throughout the CNS and are specifically well-positioned at sites of first contact with extravasating peripheral immune cells^46,47^. Their cell bodies line the parenchymal surface at the glia limitans superficialis (GLS)^46^ which contacts the immune cell-rich meningeal compartment, and their endfeet line perivascular spaces. While prior work has suggested that astrocytes can acquire features of atypical antigen-presenting cells^10,13^, here we use a pure astrocyte culture system devoid of immune cells^20^ to define the cytokine combinations sufficient to induce key components of this program. Rather than asking only whether astrocytes can express MHC-II-associated markers, we define the cytokine combinations sufficient to induce a broader T cell-interacting program, test whether stimulated astrocytes can support antigen-dependent responses of previously activated CD4+ T cells in a pure coculture system, and examine the distribution of MHC-II+GFAP+ astrocytes across inflammatory contexts in mouse and human tissue. Using OT-II and 2D2 T cell receptor transgenic systems, we test these interactions with model and CSF-relevant antigens. In parallel, we examine the presence of brain border-associated MHC-II+GFAP+ astrocytes in multiple contexts of neuroinflammation. These results place astrocytes at a critical intersection of innate and adaptive immunity in the inflamed CNS.

## RESULTS

### Interferon-responsive reactive astrocytes express antigen-presentation-associated transcriptional signatures

We recently published a large single cell RNA sequencing (scRNAseq) dataset from mouse cortex following acute sepsis modeled with i.p. injection of lipopolysaccharide (LPS). We reported a transcriptomically-defined substate of reactive astrocytes that we termed interferon-responsive reactive astrocytes (IRRAs) as they upregulated many interferon response genes. We decided to revisit this original scRNAseq dataset to further understand the potential functional output of IRRAs. As previously reported, there are multiple populations of LPS-responsive, transcriptionally distinct reactive astrocyte signatures (Figure 1A). Gene module scoring of interferon-stimulated genes (e.g. *Igtp, Ifit3, Iigp1, Isg15, Cxcl10*) shows specificity of this signature to Cluster 8. This signature overlaps with an MHC-II antigen presentation gene module (*Psmb8, Psmb9, H2-Ab1, H2-Aa, H2-Q4, H2-DMa*) in cluster 8 (Figure 1A). Dot plots of antigen presentation-related genes show the specificity of these signatures to *Igtp*+ IRRAs and *Myoc*+ border clusters 8 and 6 respectively (Figure 1B). We validated that these cells are indeed astrocytes as they lack expression of marker genes associated with microglia (*Cx3cr1, P2yr12, Tmem119*), oligodendrocyte lineage cells (*Plp1, Olig2, Mog*), vascular cells (*Flt1, Pecam1, Cldn5*), and neurons (*Snap25, Rbfox3, Dcx*), (Figure S1).

**Figure 1.**
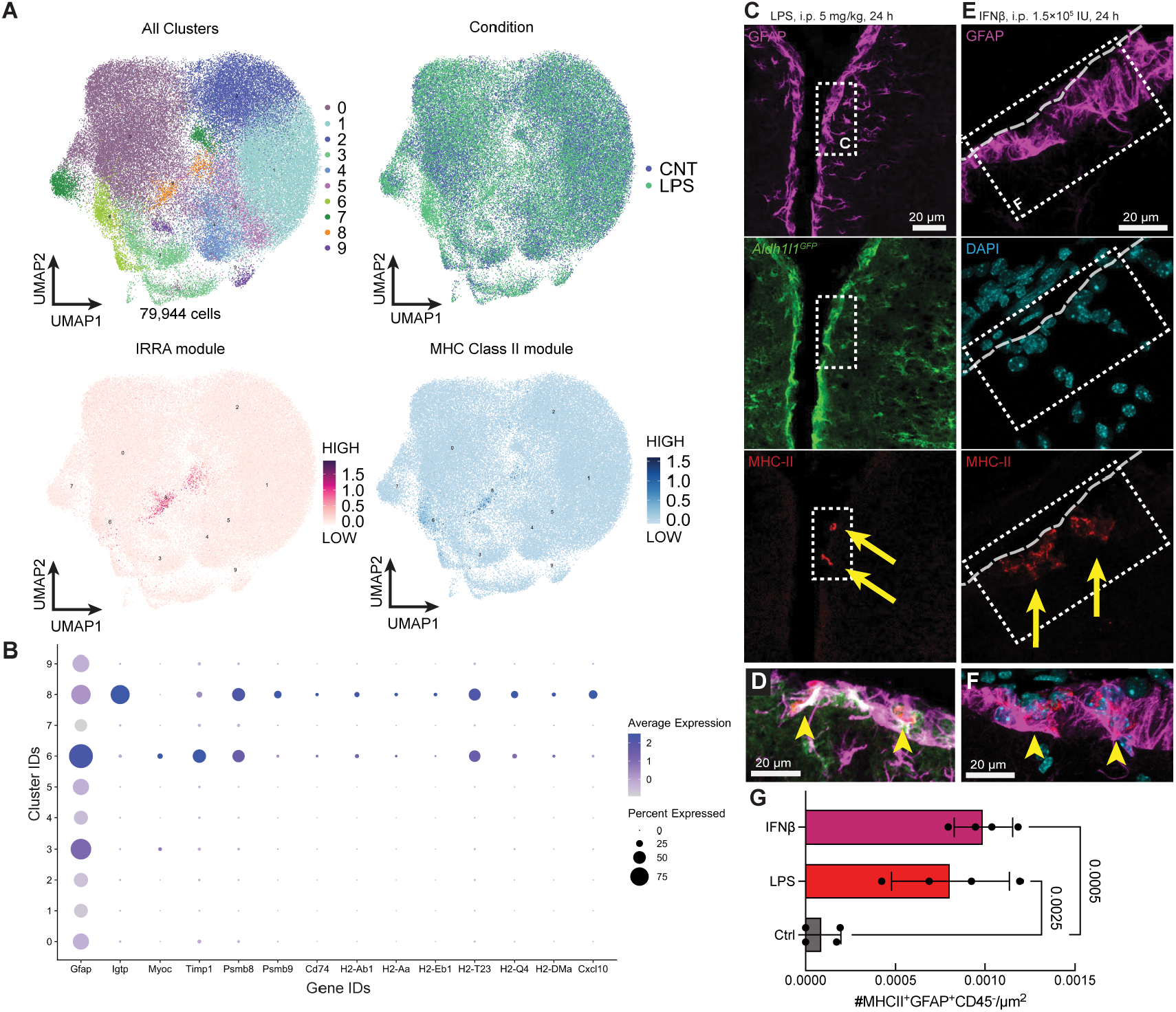
Astrocytes express antigen presentation machinery components following systemic inflammation. A. Single cell RNA sequencing (scRNAseq) data from adult mouse cortex 24 h following i.p. LPS endotoxin injection. Numerous sub-states of reactive astrocytes are present (reported elsewhere^4^). A gene module feature plot for Interferon Responsive Reactive Astrocytes (IRRAs) and genes associated with MHC Class II components occupy the same location in UMAP space. **B**. Dot plot of genes relevant to antigen presentation show specificity of transcriptional profile to clusters of interest. Gene expression is shown across clusters represented by percent expressed within each cluster and the average expression level within the cluster. **C**. Representative images following systemic injection of astrocyte reporter mice (*Aldh1l1*^eGFP^) with the bacterial cell wall endotoxin LPS (5 mg/kg, i.p. 24 h). **D**. The surface of the brain houses astrocytes (*Aldh1l1*^eGFP^ (reporter) green and GFAP (immunofluorescence) magenta) that are also positive for MHC-II (red). **E**. Representative micrograph of surface astrocytes (GFAP, magenta; *Aldh1l1*^eGFP^, green) co-labelling for MHC-II (green). Image collected 24 h following i.p. injection with IFNβ (1.5×105 IU). Like following systemic inflammation induced by i.p. LPS (A), MHC-II positive astrocytes are found in the glia limitans superficialis on the surface of the brain. **F**. (higher magnification of box in D). **G**. Quantification of MHC-II+GFAP+CD45-cell numbers normalized to tissue area. (stats = one-way ANOVA N = 4 biological replicates averaged across contralateral hemispheres). Abbreviations: CNT – control; IRRA – interferon responsive reactive astrocyte; LPS – lipopolysaccharide. Scale bars: 20 µm for all.

Here, to visualize the localization of these cells in the context of the same peripheral inflammatory insult, LPS was injected intra-peritoneally (i.p.) into adult pan-astrocyte reporter *Aldh1l1*^eGFP^ mice. *Aldh1l1*^eGFP^+MHC-II+GFAP+ astrocytes are seen along the border of the cortex near the central fissure 24 hours after peripheral LPS stimulation (Figure 1C,D), but not in saline injected animals (Figure S2A). Similar localization of MHC-II+GFAP+ astrocytes along the border of the cortex is seen after 24 hours of i.p. IFNβ (type 1 interferon) stimulation modeling systemic autoimmune conditions or persistent viral infection (Figure 1E,F), but not in saline injected animals (Figure S2B). Quantification of MHC-II+GFAP+CD45-cells (representative distinction shown in Figure S1C) reveals a significant increase in both LPS and IFNβ injected animals (Figure 1G). In both cases, these MHC-II+GFAP+ astrocytes were restricted to the cortex border and were distinct from CD45+ immune cells and leptomeningeal cells.

These findings indicate that transcriptomically distinct interferon-responsive and border-associated astrocyte populations acquire transcriptional and protein features associated with antigen presentation during peripheral inflammatory insult. In vivo, these MHC-II+GFAP+ astrocytes are localized to neuroimmune border regions that are sites of peripheral immune contact. However, these in vivo observations are descriptive – next, we aim to establish that astrocytes are sufficient for antigen presentation to T cells.

### IFNγ and proinflammatory cytokines induce the production of antigen presentation machinery

We were next interested in the signaling molecules required to induce antigen presentation machinery and MHC-II production, so we turned to an in vitro system. Pure mouse primary astrocytes were generated using a protocol adapted from Clayton et al.^20^ (Figure 2A): astrocytes were isolated from neonatal cortices and cultured in serum-free conditions with growth factors to enhance astrocyte survival over other CNS cell types. Bulk RNA sequencing of cultured astrocytes at experimentally relevant timepoints shows purity of these cultures as astrocytic markers are enriched and other cell type marker expression is minimal (Figure 2B).

**Figure 2.**
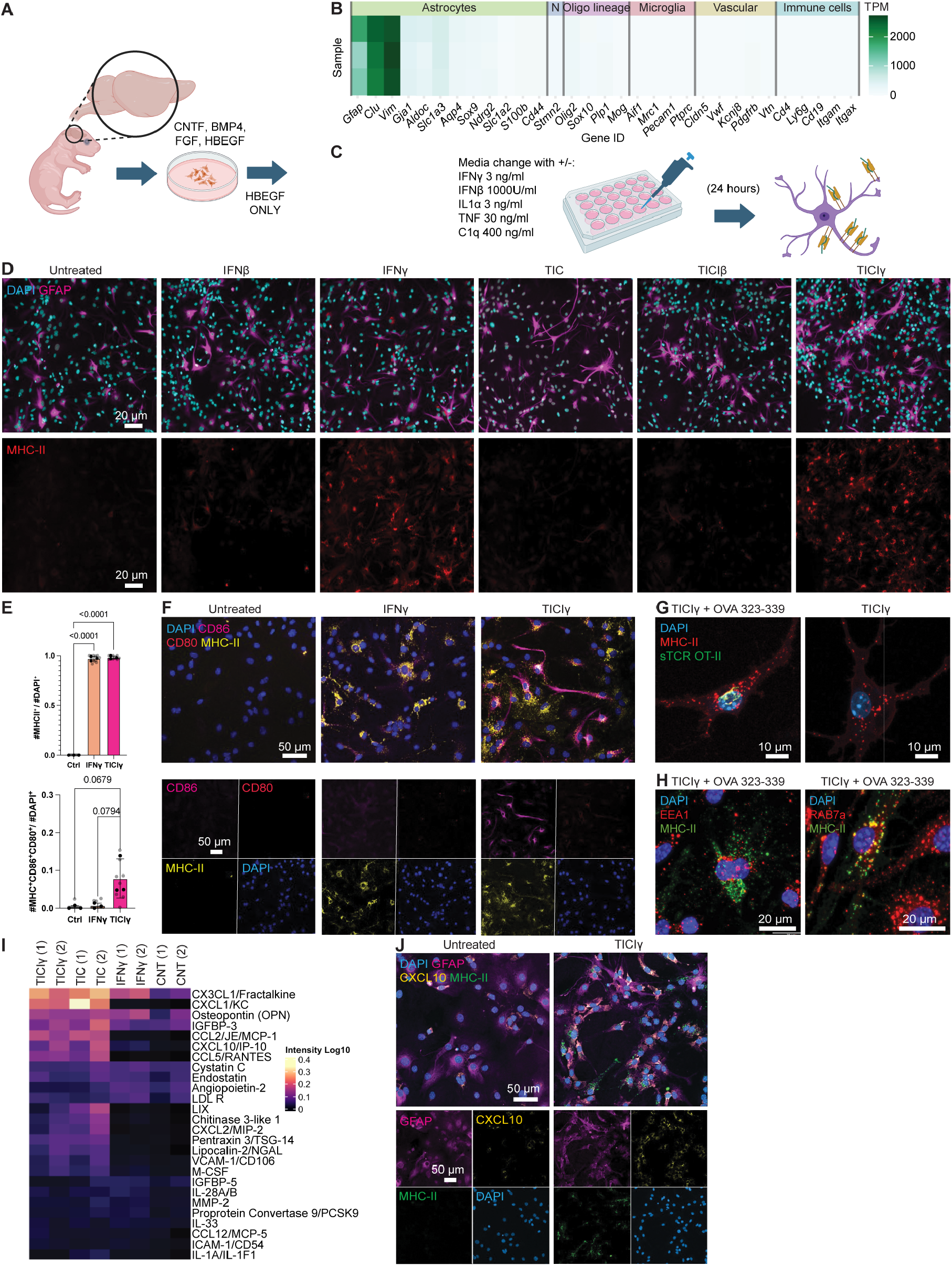
A specific combination of cytokines and interferons is required for astrocytes to express a complement of antigen presentation machinery. A. Experimental setup for isolation of postnatal mouse cortical astrocytes (using the method of Clayton et al., 2024^20^). Astrocytes were maintained in serum-free conditions, supplemented with HBEGF. **B**. RNAseq validation of cell culture purity. There is no observable expression of genes associated with neurons, oligodendrocyte lineage cells, immune cells, or vascular associate cells (endothelial cells or pericytes). Each row is an independent isolation and culture of astrocytes (N = 3). **C**. Experimental setup of induction of antigen presenting phenotype in astrocytes. **D**. Representative immunofluorescence images of astrocytes highlight that MHC-II is induced in astrocytes following treatment with interferon (IFN). Type II IFN induces MHC-II to a greater level than type I interferon. **E**. Quantification of MHC-II+ and MHC-II+CD86+CD80+ cell numbers normalized to total DAPI nuclei. (stats = one-way ANOVA N = 3 biological replicates (black dots) averaged across 3 technical replicates (gray dots)). **F**. Representative immunofluorescence images of astrocytes highlight that antigen presentation accessory proteins CD80/86 are not induced by IFNγ alone, and require the combination of TNF, IL1α, C1q, and IFNγ (TICIγ). **G**. Representative immunofluorescence images show colocalization of MHC-II molecules with fluorescently-tagged T cell receptor molecules specific to MHC-II loaded with chicken ovalbumin-derived peptide (residues 257 to 280) (top). Immunoreactivity with T cell receptor molecules is not seen when astrocytes are not treated with OVA peptide and just TICIγ. **H**. Representative immunofluorescence images show colocalization of MHC-II molecules and late endosomal marker RAB7a but not early endosomal marker EEA1 (bottom) after cytokine and OVA treatment. **I**. Heatmap of control, interferon, TIC, and TICIγ treated astrocyte secreta cytokine array. **J**. Representative immunofluorescence images showing that lymphocyte chemoattractant CXCL10 is highly produced after TICIγ treatment in GFAP+ MHC-II+ cultured astrocytes. Abbreviations: CNT – control; IFN – interferon; TPM – transcripts per million; N – neurons; Oligo – oligodendrocyte; TIC – TNF, IL1α, C1q; TICIγ – TNF, IL1α, C1q, IFNγ. Parts of this figure created with BioRender.com.

The IRRA gene signature can be induced in primary rodent astrocytes by 24 hour treatment with IFNβ (type 1 interferon) or IFNγ (type 2 interferon) +/-microglial-derived proinflammatory factors TNF, IL1α, and C1q (TIC)^4^, but whether this would also induce antigen presentation machinery was unknown. To test how these proinflammatory cues and type 1 and 2 interferons affect MHC-II production, astrocytes were treated with either IFNβ or IFNγ alone, or in combination with TIC (TICIβ or TICIγ) (Figure 2C). We observed significant MHC-II protein induction following either IFNγ or TICIγ treatment, low MHC-II positive staining following IFNβ treatment, and no appreciable induction with TIC alone (Figure 2D,E). Given the requirement for costimulation in enhancing T cell activation during antigen presentation^48,49^, we next sought to identify whether the costimulatory molecules CD80 and CD86 were present across treatment conditions. TIC alone was sufficient to induce the production of CD86 and CD80. Moreover, TICIγ stimulation led to the production of both MHC-II and costimulatory molecules (Figure 2E,F). IFNγ alone, while sufficient for robust MHC-II induction, requires the cooperation of proinflammatory TIC cues to induce a broader antigen-presentation-associated phenotype.

To assess whether astrocyte surface MHC-II could display cognate peptide for T cell receptor (TCR) recognition, we loaded cytokine-stimulated astrocytes with OVA 323-339 peptide for 24 hours and stained unpermeabilized cells with GFP-conjugated soluble T cell receptors specific to OVA-loaded MHC-II molecules, I-A^d^. Astrocytes treated under these conditions showed antigen-specific TCR staining, indicating that cytokine-stimulated astrocytes express surface MHC-II capable of displaying cognate peptide to matching TCRs (Figure 2G). Following TICIγ stimulation and OVA 323-339 loading, MHC-II protein colocalized with the late endosomal marker RAB7a but not the early endosome marker EEA1, consistent with localization of MHC-II within compartments associated with peptide loading (Figure 2H). Together, these data indicate that the combination of TIC and IFNγ induces not only MHC-II, but also additional cellular features associated with competent peptide presentation.

We then asked whether treated astrocytes produce chemokines relevant to lymphocyte recruitment. Conditioned media profiling using a multiplex Luminex cytokine array showed that lymphocyte chemoattractant chemokines, including CXCL10, CCL5, and CCL2, were strongly induced in response to TICIγ and TIC treatment but not to IFNγ treatment alone (Figure 2I). Consistent with this, immunostaining revealed CXCL10 production after TICIγ treatment (Figure 2J). These data identify a combinatorial signaling logic in astrocytes: IFNγ is sufficient for robust MHC-II induction, while proinflammatory TIC cues are required to engage accessory programs relevant to T cell interaction, including costimulation and chemokine production. Thus, in a pure primary astrocyte culture system, a combinatorial inflammatory signal involving IFNγ together with proinflammatory TIC cues is required to induce the broader antigen-presentation associated, T cell interacting phenotype observed here.

### Cytokine-stimulated astrocytes support antigen-dependent CD4+ T cell expansion in vitro

To test whether cytokine-stimulated astrocytes could support antigen-dependent restimulation of antigen-specific CD4+ T cells, we used the OT-II system, in which transgenic T cells recognize OVA 323-339 presented on I-A^d^. Astrocytes were treated with interferon or TICIγ cytokines with or without OVA peptide for 24 hours, after which media was removed and cultures were washed (Figure 3A). In parallel, CD4+CD25-T cells were isolated from OT-II or wild-type spleens and expanded under serum-free conditions with IL-2 and CD3/CD28 activator beads prior to CFSE labeling and coculture with astrocytes. This experimental design models restimulation of previously activated effector-like CD4+ T cells rather than initial priming of naïve T cells. This more closely models the trajectory of a splenic naïve T cell that is first activated in peripheral lymphoid tissues, acquires an effector phenotype, and later encounters the CNS via the circulation.

**Figure 3.**
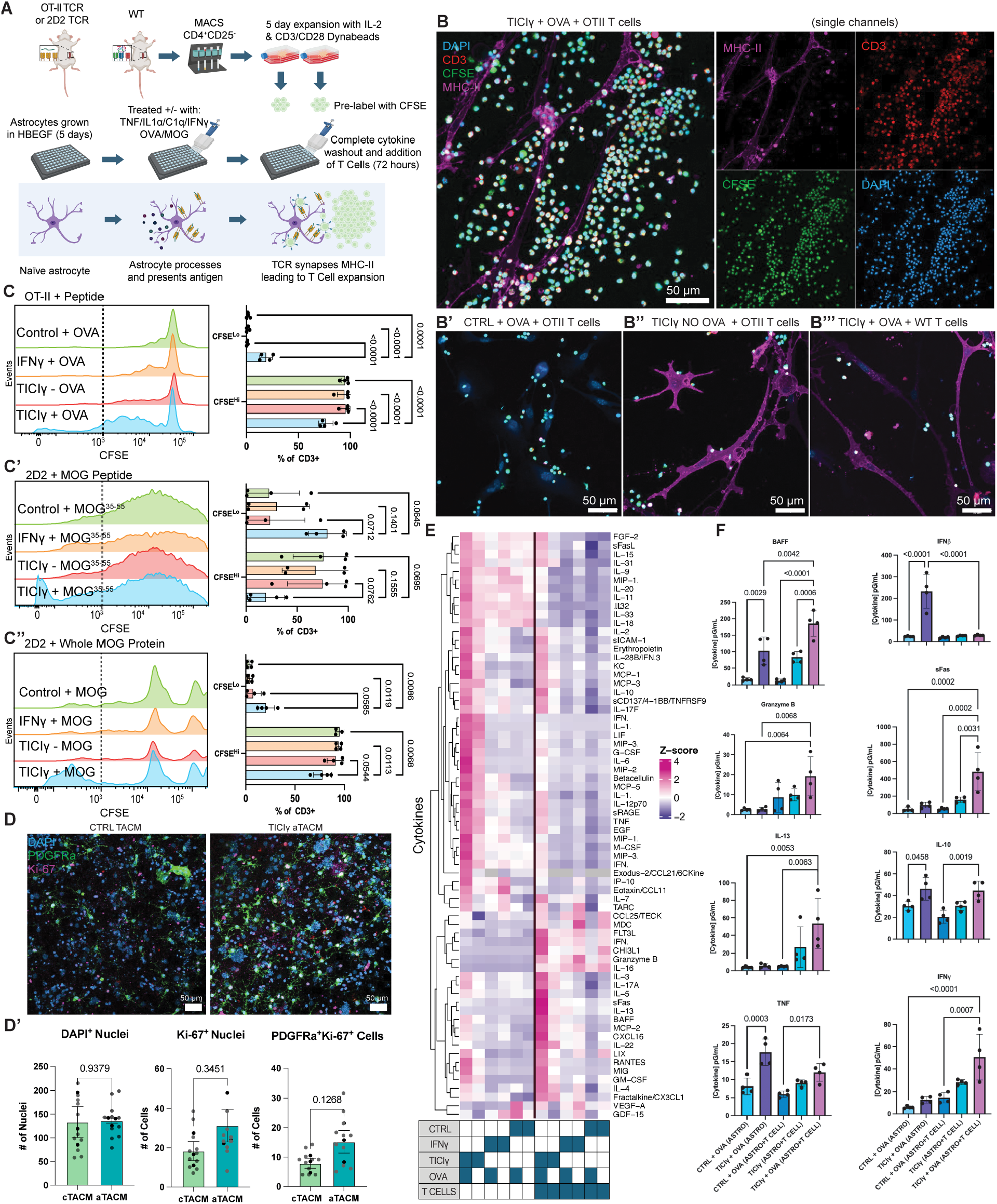
Astrocytes that express antigen presenting machinery are functional and can expand T cells. A. Depiction of experimental setup using OT-II mice to evaluate the ability of astrocytes to functionally present antigen and expand T cells. **B**. Representative immunofluorescence images of pretreated astrocytes cocultured with MACS-purified CFSE-labeled T cells. Surface staining for T cell marker CD3 and astrocyte MHC-II production show that the expansion of T cells is not seen in the absence of TICIγ pre-treatment (B’), without OVA (B’’), or with wild type T cells (B’’’). **C**. Flow cytometry of gated live CD3+ T cells after coculture. Expansion of CD3+ CFSE^lo^ T cells is seen after TICIγ and antigen pretreatment as compared to when no cytokines or antigens are applied. Single-parameter histograms of pre-labeled proliferation marker CFSE events in CD3+ gated T cells. Histograms show increased CFSE^lo^ populations (≤103) as CFSE dye is diluted during proliferation in antigen and cognate T cell combinations of OVA peptide with OT-II T cells (C), MOG35-55 peptide with 2D2 T cells (C’), and whole MOG protein with 2D2 T cells (C’’). Minimal CFSE^lo^ populations are seen in control conditions. Quantification plots at right show an increase in CFSE^lo^ populations (light blue bar) highlighting dilution of CFSE due to T cell proliferation. N = 4 bioreps per condition. Statistics are one-way ANOVA. **D**. Representative imaging of mixed rat cortical cultures after TACM application. Proliferative OPCs labeled by PDGFRa (green), and Ki-67 (magenta). **D’**. Quantification of DAPI+ nuclei, Ki-67+ cells, and double positive PDGFRa+Ki-67+ cells across activated and control conditions. Biological N = 3, with N = 4 technical replicates each. Stats are unpaired T test. **E**. Heatmap of Z scoring of cytokine concentration changes across row average per cytokine. Cytokine measured from media of untreated versus astrocytes with and without expanded T cells. Conditions are denoted by grid underneath heatmap. Each column is an average of N = 4 independent purification and culture experiments. **F**. Bar plotting of cytokine concentrations across conditions. (stats = one-way ANOVA across conditions N = 4 per condition). Abbreviations: CFSE – carboxyfluorescein succinimidyl ester, CTRL – control; IFNγ – interferon γ; OVA – chicken ovalbumin peptide; TICIγ – TNF, IL1α, C1q, IFNγ; WT – wild type. Parts of this figure created with BioRender.com.

Flow cytometry of live singlet T cells marked by CD3 shows 96% purity after MACS isolation, gated on fluorescence minus one negative stained controls (Figure S3A). When CD4+CD25-OT-II T cells were added to TICIγ+OVA treated astrocytes, we observed marked accumulation and expansion of CD3+CFSE+ T cells along the processes of astrocytes by imaging (Figure 3B). This effect was not seen in the absence of cytokine pretreatment, in the absence of OVA peptide, or when wild-type polyclonal T cells were used instead of OT-II T cells (Figure 3B’-3B’’’). To quantitatively validate these imaging observations, we performed flow cytometry on astrocyte-T cell cocultures. After gating live CD3+ T cells (Figure S3B), CFSE dilution revealed a significant increase in proliferating CFSE^lo^ populations specifically in the TICIγ+OVA condition, whereas little to no CFSE dilution was observed in the absence of OVA, TIC, or cognate TCR specificity (Figure 3C, S3B). Together, these data demonstrate that cytokine-stimulated astrocytes can support antigen-dependent expansion of previously activated CD4+ T cells in vitro. Because the responding T cells were pre-activated before coculture, these experiments model local restimulation/ expansion of effector-like CD4+ T cells rather than de novo priming of naïve T cells.

To test a more physiologically relevant CNS antigen, we repeated this coculture paradigm using transgenic CD4+ T cells from 2D2 TCRMOG mice, which recognize MOG-derived antigen. After coculture with TICIγ-stimulated astrocytes, we again performed flow cytometry, measuring CFSE dilution in live CD3+ T cells, and observed expansion selectively in the TICIγ+MOG35-55 peptide (Figure 3C’, S3B’) and TICIγ+whole MOG protein conditions (Figure 3C’’, S3B’’). These findings indicate that stimulated astrocytes can not only present exogenously supplied cognate peptide, but can also support T cell responses after exposure to intact MOG protein, consistent with the capacity to take up and process whole protein antigen into a stimulatory form. This whole-protein experiment is important because, unlike peptide-loading assays, it requires antigen uptake and intracellular processing before productive presentation to 2D2 T cells.

As professional antigen presenting cells have the capacity to expand naïve splenic T cells, we repeated these experiments without the initial expansion and stimulation of T cells in culture. Flow cytometry of naïve T cell-astrocyte cultures reveals a large amount of cell death (16.5%-60.9%) across conditions including TICIγ+OVA and the absence of OVA or cytokines. In coculture, these unstimulated T cells are likely more susceptible to death given a possible absence of cytokine signaling to maintain them whereas effector cells may be more resilient in culture with astrocytes^50-52^. Interestingly, there still was a higher proportion of CD3+ T cells in the TICIγ +OVA (71.4%) as compared to controls (27.9% - 36.3%) suggesting either a greater survival or limited expansion in the context of stimulated astrocytes. Across all conditions, the CFSE^hi^ population is largely absent, whereas a population of CFSE^lo^ T cells was observed (Figure S3C). This may reflect naïve T cells undergoing limited proliferation but failing to survive in the absence of appropriate supportive factors^52^.

These observations suggest that, under the present culture conditions, astrocytes are more effective at supporting antigen-dependent responses of previously activated effector-like T cells than at sustaining naïve T cells. Given that the CNS is more likely to encounter infiltrating antigen-experienced effector T cells than naïve T cells, we focused our interpretation on astrocyte-mediated restimulation rather than naïve T cell priming.

To further understand the downstream functions of astrocyte-activated effector T cells, we added the T cell-Astrocyte Conditioned Medium (TACM) from these experiments to mixed rat cortical cultures. Addition of activated TACM (after TICIγ+OVA treatment of astrocytes and T cell coculture) caused a noticeable increase in PDGFRa+ cells by immunostaining as compared to control TACM (absence of astrocyte cytokine stimulation) (Figure 3D). Quantification of these immunostained cultures revealed no increase in total DAPI+ nuclei and Ki-67+, but did show an increase in double positive PDGFRa+Ki-67+ cells (Figure 3D’) after activated TACM treatment as compared to control TACM. These data identify a potential downstream consequence of astrocyte–T cell crosstalk in vitro, whereby conditioned medium from activated cocultures promotes OPC proliferation markers in mixed cortical cultures. To probe the T cell phenotypes downstream of astrocyte mediated activation and expansion, we profiled the concentration of 68 cytokines commonly involved in T cell mediated immune function in all conditioned ACM and TACM. Z scoring and hierarchical clustering of cytokine production revealed a marked shift in activated TACM composition, consistent with heterogenous T cell responses in the same condition (Figure 3E, S4A). Interestingly many cytokines produced at baseline by activated astrocytes are absent unless there is sufficient T cell expansion and stimulation to maintain this activation in vitro. Quantification of cytokine concentrations shows no delineation for the expansion of one T cell subtype rather, the activation of multiple pathways that modulate several lymphoid cells and substates such as: B cell activation through BAFF, propagation of type II IFN or Th1 responses including a shift away from IFNβ-associated signaling and toward IFNγ- and TNF-associated inflammatory outputs. Alternatively, Th2, Treg, and cytotoxic T responses may be present with the presence of IL-13, IL-10, and sFas/ Granzyme B, respectively (Figure 3F). Taken together, these results suggest that astrocyte-mediated T cell restimulation in vitro can generate heterogeneous cytokine outputs with downstream effects on other CNS cell populations, although the precise pathways and in vivo relevance remain to be determined. These findings identify a potential downstream consequence of astrocyte-T cell crosstalk in vitro, but the mediators and in vivo relevance of this effect remain unresolved.

### Human iPSC-derived astrocytes upregulate antigen-presentation-associated machinery in response to inflammatory cytokine stimulation

We next asked whether human astrocytes in a multicellular iPSC-derived CNS culture can upregulate antigen-presentation-associated machinery in response to inflammatory cytokine stimulation. To do so we used human iPSC-derived CNS multicellular cultures derived from reprogrammed PBMCs from six independent donors^53^ (Figure 4A). Cell cultures, including neurons, oligodendrocytes, OPCs, astrocytes, and microglia, were treated with a vehicle, TNF, or IFNγ (for a total of 18 distinct samples). We applied high-plex probe-based single cell transcriptomics using the 10x Flex platform, encompassing probes for 18,129 genes. The resulting dataset revealed clearly defined distinct cell types, each exhibiting transcriptional responses to cytokine stimulation (Figure 4B,C). Quality control of culture sequencing showed that clusters are not driven by individual donor patient profiles, mitochondrial count, or unique molecular identifiers (Figure S5A).

**Figure 4.**
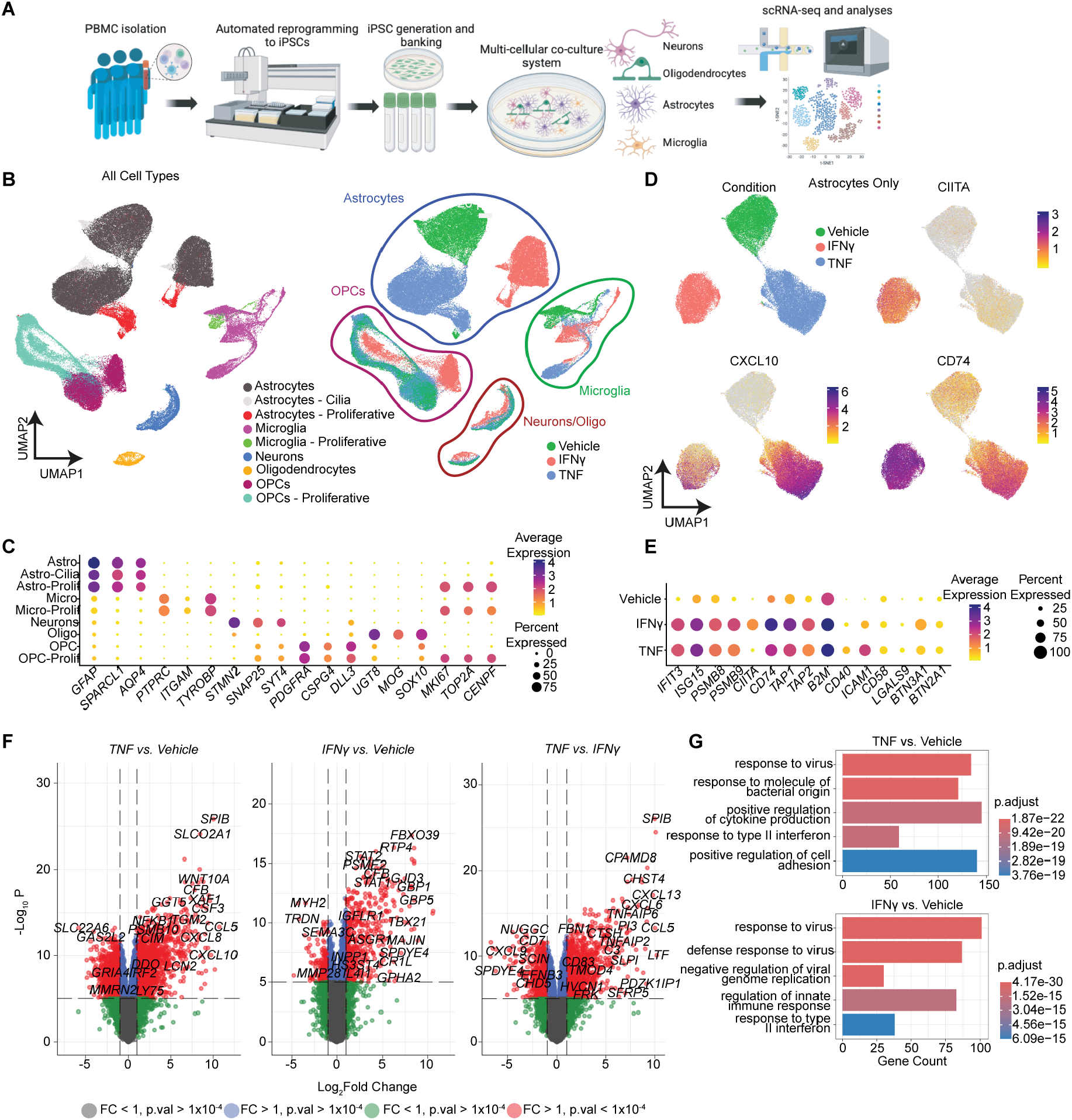
Human iPSC astrocytes express AP machinery in response to cytokine stimulation. A.Schematic of PBMC collection, iPSC reprogramming, differentiation to CNS cells, and sequencing. See Table S1 for donor demographics. **B**. UMAP of all sequenced cell types split by cell type and condition. N = 6 individual patient iPSC lines were differentiated (see Figure S5A), cultured, treated, and sequenced. **C**. Dot Plot of marker gene expression across clustered cell types. **D**. UMAP of Astrocytes only split by condition and feature plots of AP machinery genes *CIITA, CXCL10*, and *CD74*. **E**. Dot plot of antigen presentation and T cell communication relevant gene expression across conditions in astrocytes. **F**. Volcano plots of differentially expressed genes between conditions. Upregulated genes correspond to first condition listed in title i.e. TNF (+ L_2_FC) vs Vehicle (-L_2_FC). **G**. GO term gene analysis of top 5 pathways upregulated colored by p value. Parts of this figure created with BioRender.com.

Because the targeted human probe set did not include polymorphic MHC genes, these experiments do not directly measure human MHC-II transcripts; rather, they support induction of a broader antigen-presentation-associated program through *CIITA, CD74*, antigen-processing genes, and T cell-interaction genes. While only microglia showed appreciable expression of *CIITA*, or the MHC Class II transactivator, at baseline, it was robustly upregulated in astrocytes (and OPCs) following IFNγ stimulation (Figure 4D,E, Figure S5B,B’). Feature plots of subsetted astrocytes revealed increased *CIITA* expression in response to IFNγ. Expression of *CXCL10* and *CD74*, an MHC class II chaperone, was also increased following cytokine treatment. *CXCL10*, a lymphocyte chemoattractant chemokine, was more potently up-regulated by TNF than by IFNγ, whereas CD74 was more strongly induced by IFNγ than by TNF (Figure 4D). These effects were not driven by donor variability (Figure S5C). Increases in expression of additional necessary machinery for antigen processing (*PSMB8, PSMB9, TAP1, TAP2*), MHC-I (*B2M*), and T cell communication/costimulation (*ICAM1, BT-N3A1, BTN2A1, LGALS9*) are also seen in astrocytes after both TNF and IFNγ treatment (Figure 4E). Differential gene expression analysis of pseudobulked subsetted astrocytes comparing treatment conditions reveals the upregulation of cytokine transcription in the TNF-treated condition, whereas IFNγ-treated condition shows an increase in interferon-stimulated genes, as expected (Figure 4F). Gene ontology analyses of these differentially expressed genes in TNF and IFNγ conditions show that the top 5 pathways upregulated in astrocytes include those relevant to responding to pathogenic insult or cytokine release (Figure 4G).

Further analysis of subsetted astrocytes across these conditions reveals broad reactive astrocyte marker transcription that is increased in response to TNF and IFNγ (Figure S5D). Baseline astrocyte reactivity across conditions likely reflects microglial production of cytokines in culture (Figure S5E). Astrocytes also express lymphocyte attractant chemokine genes in response to treatment (*CXCL10, CCL2, CCL5*) (Figure S5D) similar to those seen in mouse astrocytes after treatment (Figure 2I,J). Subsetted microglia from these cultures also express genes corresponding to immune-modulatory cytokines, especially in response to IFNγ stimulation (Figure S5E). Interestingly, TNF addition reduces the expression of a multitude of these cytokine genes, a potential indication of either IFNγ potency in stimulating microglia or a shift in inflammatory transcriptional profile away from proinflammation. OPCs subsetted from these cultures show only moderate increases in proliferative markers (*MKI-67, TOP2A*) in response to TNF stimulation, and no change in lineage commitment-related genes (Figure S5F). These data suggest that additional signals, potentially including T cell-derived cues, may be required to induce stronger proliferative responses in OPC populations. Together, these human iPSC-derived CNS culture data support the conclusion that inflammatory cytokine exposure induces human astrocytes to upregulate antigen-presentation-associated and lymphocyte-interacting machinery. In this multicellular context, IFNγ stimulation occurs in the presence of microglia and other potential sources of proinflammatory cues, which likely contributes to the broader astrocyte responses observed here. These findings also suggest that, in the absence of microglia and associated proinflammatory cues, astrocytes may upregulate only a partial antigen-presentation program which includes MHC-II induction, but less robust accessory machinery and reduced T cell-supporting function, as suggested by recent work^54^. These findings therefore support conservation of an inducible astrocyte immune-interaction program in human cells, while also underscoring the importance of cellular context in shaping its full expression.

### MHC-II+GFAP+ astrocytes are present across multiple neuroinflammatory contexts

Reactive astrocytes and interferon activation are ubiquitously present in many disease contexts. We therefore decided to test whether MHC-II+GFAP+ astrocytes are present in several animal models of inflammation and disease that have previously been shown to involve reactive astrocytes, interferon responses, and T cells. We chose experimental autoimmune encephalomyelitis (EAE, a mouse model of demyelination) and viral infection with SARS-Co-V-2 as both induce a strong peripheral inflammatory response.

EAE, a mouse model of MS, is an autoimmune neuroinflammatory disease characterized by increased MHC-II and systemic IFN production as well as infiltration of autoreactive CD4+ T cells into the CNS. This results in lesion pathology, as T cells attack cognate myelin antigens and exacerbate local immune response and cell death. We therefore induced MOG35-55 EAE and examined the localization of MHC-II expressing astrocytes. Effectively, EAE induced mice developed spinal cord lesions marked by T cell and MHC-II+ cell accumulation in the spinal cord (Figure S6A,B). Mice reached peak EAE score 3 between days 13 and 15 and showed a consistent ∼17% reduction in body weight (Figure S6B). At peak EAE, MHC-II+GFAP+ astrocytes were significantly increased at the cortex border while sparsely found in MOG-controls (Figure 5A-D). These MHC-II+ astrocytes were situated near CD3+ T cells (Figure 5B,C). These data demonstrate the emergence of MHC-II+GFAP+ astrocytes at the brain border in EAE and their close spatial association with CD3+ T cells, supporting the hypothesis that astrocytes may participate in local neuroimmune interactions during autoimmune inflammation.

**Figure 5.**
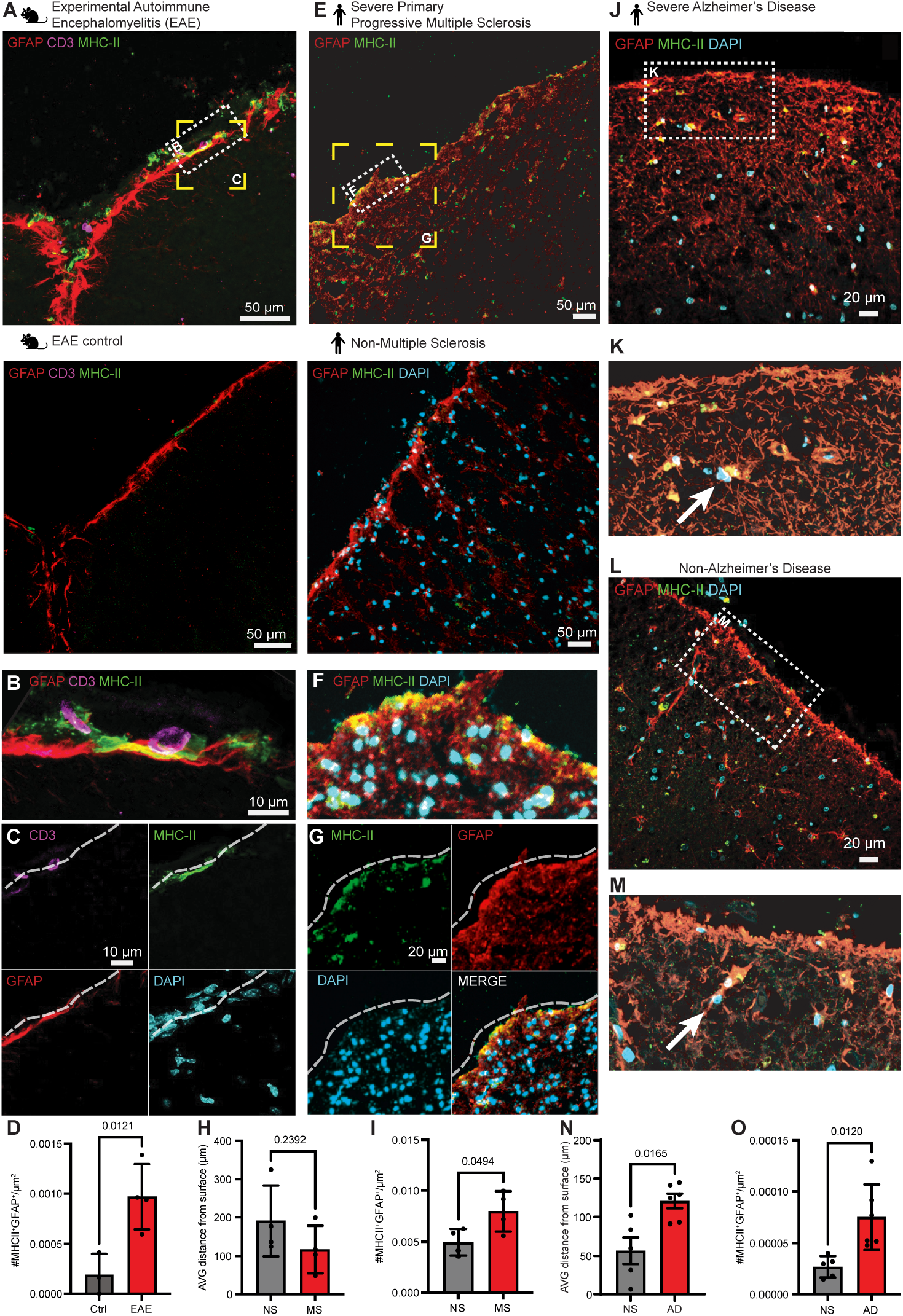
In vivo examples of surface astrocytes expressing MHC-II. A. Representative micrograph of surface astrocytes (GFAP, red) co-labelling for MHC-II (green), in close association with CD3+ T cells (magenta). Image collected EAE peak 19 days following induction of EAE (top) or in MOG-EAE controls (bottom) in adult female mice. CD3+ T cells do not enter the brain and interact with astrocytes, inducing MHC-II in the glia limitans superficialis on the surface of the brain. **B**. (higher magnification of box in A). **C**. (higher magnification of box in A). Light dashed line shows surface of brain. **D**. Quantification of MHC-II+GFAP+ cell numbers normalized to tissue area. (stats = unpaired T test N = 4 biological replicates averaged across contralateral hemispheres). **E**. Representative micrograph of human surface astrocytes (GFAP, red) co-labelling for MHC-II (green). Image collected in human pre-frontal cortex during severe primary progressive multiple sclerosis (top) or in non-symptomatic controls (bottom). (Table S2). **F**. (higher magnification of box in G). **G**. (higher magnification of box in G). Light dashed line shows surface of brain. **H**. Quantification of MHC-II+GFAP+ astrocyte distance to brain surface normalized to tissue area. (stats = unpaired t test N = 4 MS 4 NS, averaged over 3 imaging ROIs/sample). **I**. Quantification of MHC-II+GFAP+ astrocyte numbers normalized to tissue area. (stats = unpaired t test N = 4 MS 4 NS, averaged over 3 imaging ROIs/ sample). **J**. Representative micrograph of human surface astrocytes (GFAP, red) co-labelling for MHC-II (green). Image collected in human pre-frontal cortex during severe Alzheimer’s Disease. **K**. (higher magnification of box in I). **L**. Representative micrograph of human surface astrocytes (GFAP, red) co-labelling for MHC-II (green). Image collected in human pre-frontal cortex in non-symptomatic controls (right). **M**. (higher magnification of box in K). **N**. Quantification of MHC-II+GFAP+ astrocyte distance to brain surface normalized to tissue area. (stats = unpaired t test N = 6 AD 5 NS, averaged over 3 imaging ROIs/sample) (Table S2). **O**. Quantification of MHC-II+GFAP+ astrocyte numbers normalized to tissue area. (stats = unpaired t test N = 6 AD 5 NS, averaged over 3 imaging ROIs/sample) (Table S2).

As EAE was able to induce MHC-II+ astrocytes, likely due to peripheral inflammation and strong IFN signaling associated with this model, we next investigated whether a viral infection (which also induces robust IFN signaling peripherally) would also induce this subset of astrocytes. The localized sequestration of pathogens at brain border interfaces plays a key role in protecting the CNS from infection^55^. As antigen processing and presentation are robust means of communicating viral infection to the adaptive immune system, we wanted to interrogate MHC-II+ astrocyte generation during viral infection^56^. We chose to model SARS-CoV-2 infection as it is known to result in systemic prolonged interferon responses and in some cases chronic neurological sequela57. To do so, 2,000 pfu of SARS-CoV-2 WA-1 were inoculated intranasally into K18-hACE2 mice. After 5 days of infection, MHC-II+GFAP+ astrocytes were seen in the GLS space and also colocalized to SARS-CoV-2 nucleocapsid staining (Figure S6C-E), whereas none were seen along the parenchymal surface in saline-administered mice (Figure S6D). After 3 days of IFNγ blockade, no MHC-II+GFAP+ astrocytes were seen along the parenchymal surface while viral nucleocapsid burden was heightened in the cortex (Figure S6E). Given that antigen-presentation-associated machinery induced in cytokine-stimulated astrocytes in vitro, one possibility is that these border-localized MHC-II+GFAP+ astrocytes engage immune signaling relevant to viral exposure at CNS interfaces. However, direct demonstration of astrocyte-dependent antigen presentation in vivo will require further study.

We next sought to localize MHC-II+GFAP+ astrocytes in the context of human disease. In Severe Primary Progressive Multiple Sclerosis (PPMS) MHC-II+GFAP+ astrocytes are seen (Figure 5E-G) along the border of the prefrontal cortex (Figure 5F,G) as well as some MHC-II+GFAP+ cells. Some MHC-II+GFAP+ astrocytes were observed in Non-Multiple Sclerosis brains (Figure 5E) but there was a significant increase in PPMS patient cortices (Figure 5I). MHC-II+GFAP+ astrocytes are seen at the borders of hypercellular areas likely corresponding to active lesions (Figure S6F) as well as near large vessels in the prefrontal cortex (Figure S6F’). MHC-II+GFAP+ astrocytes are also noted in the pre-frontal cortex of Severe Alzheimer’s Disease (AD) patients (Figure 5J,K) and their age-matched (∼77 y.o.) non-symptomatic controls (Figure 5L,M). Quantification of these populations reveals that there is a significant increase in total MHC-II+GFAP+ astrocytes and their distance from the parenchymal surface (Figure 5N,O). The presence of some MHC-II+GFAP+ astrocytes in non-neurological aged control tissue suggests that age and cumulative inflammatory exposure may influence baseline abundance, whereas chronic inflammatory disease states are associated with increased numbers and broader distribution of these cells.

Together, these data show that MHC-II+GFAP+ astrocytes are present at outer CNS borders across viral, autoimmune, and neurodegenerative contexts in mice and humans. Their localization places them at a neuroimmune interface well positioned for interaction with peripheral immune cells. Nevertheless, our in vivo data are descriptive and should be interpreted as establishing the presence and localization of these cells, rather than proving the magnitude of their functional contribution to antigen presentation in situ.

## DISCUSSION

Here, we show in a pure astrocyte culture system that IFNγ together with proinflammatory TIC cytokine cues cooperatively induce an antigen-presentation-associated program in astrocytes, including MHC-II, costimulatory molecules, and chemokines relevant to lymphocyte recruitment. Under these conditions, stimulated astrocytes support antigen-dependent expansion of previously activated CD4+ T cells in vitro. In parallel, single-cell transcriptomic reanalysis and tissue staining identify MHC-II+GFAP+ astrocytes at CNS borders across multiple inflammatory contexts in mice and humans. Thus, our study distinguishes between a functional in vitro demonstration of astrocyte-supported T cell restimulation and descriptive in vivo evidence that astrocytes expressing antigen-presentation-associated machinery emerge at neuroimmune interfaces during inflammation. This distinction is central to the interpretation of our study, and these findings expand prior work showing that astrocytes can express MHC-II-related molecules by defining cytokine conditions that induce a broader T cell-interacting phenotype and by demonstrating antigen-dependent CD4+ T cell expansion in a pure astrocyte coculture system.

The capacity of astrocytes to act as atypical antigen presenting cells in neuroinflammatory contexts adds mechanistic and functional detail to our understanding of their immune effector functions – positioning them as plausible contributors to local effector T cell reactivation during neuroinflammation. Consequently, the implication of this gain-of-mechanism in reactive astrocytes may vary across contexts.

### How may MHC-II+ astrocytes at CNS borders modulate immunity?

At brain border regions such as the GLS, perivascular spaces, and meningeal interfaces, astrocytes are well positioned to encounter infiltrating immune cells. Their anatomical location raises the possibility that MHC-II+ astrocytes participate in local communication with the adaptive immune system in ways that could either restrain or amplify inflammation depending on context. In autoimmune settings such as multiple sclerosis, such interactions could in principle support local restimulation of autoreactive CD4+ T cells, whereas in infection they might contribute to immune surveillance against foreign antigen. These possibilities remain hypotheses that require direct in vivo testing. In our in vitro system, conditioned medium from activated astrocyte–T cell cocultures promoted proliferation-associated OPC markers, suggesting one possible downstream consequence of astrocyte–T cell signaling. However, the molecular basis of this effect and its relevance in vivo remain unresolved, and these findings should be interpreted as hypothesis-generating. Potential mechanisms linking astrocyte-T cell signaling to OPC responses remain to be determined.

### What T cell subtypes are engaged by astrocytes?

An important open question is which CD4+ T cell states are preferentially supported by cytokine-stimulated astrocytes. In our coculture system, cytokine profiling suggested heterogeneous T cell-associated outputs rather than selective enrichment of a single canonical subtype, implying that astrocyte-mediated restimulation may be strongly shaped by local context. Future studies will be needed to define how antigen dose, costimulatory balance, inflammatory cytokines, and astrocyte state influence the differentiation or maintenance of Th1-, Th2-, Th17-, or regulatory-like programs. Further experiments should address astrocyte response heterogeneity in vitro in response to our treatment paradigm as this may influence T cell specialization. If astrocytes do in fact respond homogeneously, further experiments should also address the integrated strength, duration, and context of signaling received by individual T cells as this heterogeneity in T cell subtypes may emerge from variation in TCR affinity for peptide–MHC complexes, stochastic differences in signaling pathways, cytokine consumption, metabolic state, and prior activation history of the responding T cells. In this case, manipulation of antigen concentration, duration of stimulation, cell density, and cytokines secreted by activated T cells can generate more specific and localized signaling environments that promote distinct transcriptional differentiation programs. IFNγ-driven upregulation of MHC-II and associated machinery suggests a preferential induction of Th1-polarized cells. However, the potential for astrocytes to present antigen in a context that promotes Treg induction or maintenance also exists, particularly when astrocytic presentation is accompanied by anti-inflammatory signals such as IL-10 or TGF^58^.

### What are the temporal dynamics of this MHC-II+ phenotype?

The induction of this astrocyte immune-interaction phenotype appears tightly linked to inflammatory cues coming from other cell types, particularly IFNγ together with proinflammatory cues. In our experiments, astrocytes upregulated MHC-II associated machinery rapidly, often within 24-48 hours, in both rodent and human systems. Whether this represents a transient reactive state or a longer-lasting functional reprogramming remains unknown and will require time-course studies, withdrawal paradigms, and lineage-resolved analyses.

Our findings highlight multiple hypotheses of downstream multi-cellular interactions that could further neuroinflammation. In response to T cell expansion and activation, microglia, astrocytes, and the vasculature can respond to cytokines produced by T cells. T cells could also recognize and target specific antigens, causing cell death. As mentioned above, while other CNS cell types may have the capacity to produce antigen-presentation machinery and/or stimulate T cells, they are likely involved in the propagation of neuroinflammation of which astrocytes are poised to play a central role. A key limitation of the present study is that astrocyte-dependent antigen presentation was demonstrated functionally in vitro but not directly tested in vivo using astrocyte-specific perturbation of MHC-II or adoptive antigen-specific T cell readouts. Accordingly, our in vivo data establish the presence, localization, and disease association of MHC-II+GFAP+ astrocytes, but do not by themselves define the magnitude of astrocyte contribution to T cell activation within intact tissue.

Overall, our findings show that inflammatory cytokine combinations, particularly IFNγ together with proinflammatory cues, induce astrocytes to acquire a broader antigen-presentation-associated phenotype and to support antigen-dependent restimulation/expansion of previously activated CD4+ T cells in vitro. In parallel, MHC-II+GFAP+ astrocytes emerge at CNS borders across multiple inflammatory settings in mice and humans, placing them at a strategic neuroimmune interface. These in vivo observations are descriptive and do not establish astrocyte-specific necessity for antigen presentation in situ. We therefore view the present study as defining an inducible astrocyte immune-interaction program and a framework for future in vivo tests of its functional importance during neuroinflammation.

## METHODS

### Animals and ethics statement

All animal procedures were performed in accordance with institutional and national guidelines for the care and use of laboratory animals and were approved by the Institutional Animal Care and Use Committee (IACUC) at NYU Grossman School of Medicine. Adult C57BL/6J mice (RRID:IMSR_ JAX:000664), OT-II B6.Cg-Tg(TcraTcrb)425Cbn/J (RRID:IMSR_JAX:004194), and *Aldh1l1*^eGFP^ (Tg(Aldh1l1-EGFP)OFC789Gsat/Mmucd (RRID:IMSR_JAX:026033) of either sex were used for all experiments. Mice were housed under a 12-hour light/dark cycle with ad libitum access to food and water. For SARS-CoV2 related experiments, B6.Cg-Tg(K18-ACE2)2Prlmn/J (K18-hACE2) mice (stock no. 034860) were purchased from the Jackson Laboratory and subsequently bred and housed at Yale University. Twelve-week-old female mice were used. All procedures used in this study complied with federal guidelines and the institutional policies of the Yale School of Medicine Animal Care and Use Committee.

### In vivo inflammatory models

#### Virus stock, infection, and antibody treatment

As reported previously^59^ Vero E6 cells overexpressing angiotensin-converting enzyme 2 (ACE2) and TMPRSS2 [kindly provided by B. Graham at the National Institutes of Health Vaccine Research Center (NIH-VRC)] were infected with SARS-CoV-2 isolate hCOV-19/USA-WA1/2020 (NR-52281; BEI Resources) at a M.O.I. of 0.01 for 2 days until sufficient cytopathic effect was noted. After incubation, the supernatant was clarified by centrifugation (5 min, 500g), filtered through a 0.45-μm filter and stored at -80 °C. Viral titers were measured with a standard plaque assay by using Vero E6 cells. For virus challenge, mice were anesthetized by using 30% v/v isoflurane diluted in propylene glycol. Using a pipette, 50 μl containing 2000 plaque-forming units (PFU) SARS-CoV-2 was delivered intranasally. For IFN-γ blockade, mice were treated with 200 µg (IFN-γ, Clone H22; Biolegend) blocking antibodies. For CD4+ T cell depletion, mice were treated with 200 µg of a CD4 depletion antibody (Clone GK1.5; BioX-Cell). All antibodies were diluted in 200 µl PBS and injected intraperitoneally 2 days after virus infection, with treatment continuing for 3 days. Experiments involving SARS-CoV-2 infection were performed in a biosafety level 3 facility with approval from the Yale Institutional Animal Care and Use Committee and Yale Environmental Health and Safety.

#### LPS and IFNβ injections

Adult C57BL/6J mice (8–10 weeks old) were administered a single intraperitoneal (IP) injection of lipopolysaccharide (LPS; E. coli O111:B4, Sigma L2880) at a dose of 5 mg/ kg to induce systemic inflammation. In parallel experiments, recombinant murine interferon β (IFNβ; PeproTech) was injected IP at 1 µg/20g mouse to model systemic interferon release. Control animals received equivalent volumes of sterile PBS. Mice were sacrificed and perfused with cold PBS before immersion fixation 24 hours post-injection for brain tissue collection.

#### Experimental Autoimmune Encephalomyelitis (EAE)

EAE was induced in 8 week-old female C57BL/6J mice using the MOG 35–55 EAE induction kit (Hooke Laboratories, EK-2110), following the manufacturer’s instructions. Briefly, mice received subcutaneous injections of MOG35– 55 peptide emulsified in complete Freund’s adjuvant on day 0, followed by intraperitoneal injections of pertussis toxin on days 0 (2 hours after MOG induction) and 1. Mice were monitored daily and scored using the standardized Hooke scoring scale (0–5). Animals were euthanized at peak disease or control-matched timepoints.

### Cell culture

#### Primary astrocyte isolation and culture

Primary astrocyte isolation and culture protocol was adapted from Clayton et al^20^. Primary cortical astrocytes were isolated from postnatal day 0–2 mouse pups. Cortices were dissected in ice-cold DPBS, meninges removed, and tissue enzymatically dissociated with papain (Worthington 9001-73-4 1:300) followed by mechanical trituration. Cells were plated in 10 cm cell culture dishes coated with poly-D-lysine (Sigma 1:500) and Laminin (Gibco 1:200). Growth media consisted of DMEM/F12 (Thermo Fisher 11320033) and Neurobasal (Thermo Fisher) (1:1 DMEM:Neurobasal) supplemented with B27 (Gibco 1:50), NEAA (Gibco 1:100), N-acetylcysteine (5 ug/mL 1:1000), N2 (Gibco 1:100), HBEGF (Sigma E4643 5 ng/mL), CNTF (PeproTech 450-13 10 ng/mL), BMP4 (Stemcell 78211.1 10 ng/mL), FGF2 (Stemcell 78003 20 ng/mL), 1% penicillin-streptomycin, 30% glucose (1:100), and 2 mM L-glutamine at 37 °C with 5% CO_2_ After 2 days media was replaced without B27. After 4 more days of culture with growth factors, astrocyte monolayers were removed from dishes with TrypLe (Thermo 12-605-010) and frozen at -80 ºC with CryoStor (Stemcell 07930). When used experimentally, astrocytes were thawed and plated in wells coated with PDL and Laminin. Media consisted of the Neurobasal DMEM/F12 (Thermo Fisher 11320033) supplemented with HBEGF, N2, NEAA, 1% penicillin-streptomycin, 30% glucose, N-acetyl cysteine, and 2 mM L-glutamine at 37 °C (as above) with 5% CO_2_. Astrocytes were ultimately used at ∼80% confluency after ∼5 day culture and media was half changed every other day.

#### Dissociated rat cortical cultures

Cortical cultures were isolated from Sprague Dawley rat pups (postnatal 1-2). The cerebrum was microdissected and washed twice in ice-cold HBSS (4.2 mM NaHCO_3_, 1 mM HEPES, pH 7.35, 300 mOsm) before a 25min digestion in papain solution (1 mL HBSS + 58U papain) at 37 °C/5%CO_2_. 4 μL of DNase I (0.2 M), 0.5 mM CaCl_2_, and 1 mM MgCl_2_ were added, and the mix was inverted gently a few times. After 4 more minutes of digestion in papain at 37 °C/5%CO_2_, HBSS containing 20% FBS was added to stop digestion. The tissue was washed additional times, then triturated in 1 mL HBSS + 8 μL of DNase I (0.2M) using 3 fire polished Pasteur pipettes of decreasing diameter. The remaining cell suspension was pelleted by centrifugation (10 min at 1000 RPM, 4 °C) through a 4% BSA cushion, resuspended in trituration medium, then pelleted again (10 min at 1000 RPM, 4 °C). After resuspension in culture media (astrocyte base media (see above) +B27), the cell suspension was filtered through a 70 μm strainer. Cells were counted with a hemocytometer and plated 30,000 cells per well of 96-well plates. 1 day following dissection (DIV, days in vitro), cells were washed with PBS to remove debris, then a half media change was performed every 48 hours. On DIV7, cells were treated with T cell astrocyte conditioned medium. After 24 hours, cells were fixed with 4%PFA.

#### T cell isolation and in vitro activation

##### CD4+ CD25-T cell isolation

Spleens were harvested from adult C57BL/6J and OT-II mice. Single-cell suspensions were generated via mechanical dissociation and filtration through a 70 µm strainer. Red blood cells were lysed using eBioscience™ 1X RBC Lysis Buffer (Invitrogen 00-4333-57). CD4+CD25-T cells were isolated by negative selection using magnetic-activated cell sorting (MACS) with CD4+ T cell and CD25 isolation kits (Miltenyi Biotec 130-104-454 & 130-091-072) or Naïve isolation kit (Miltenyi Biotec 130-104-453), following the manufacturer’s protocol. T cell purity (>96%) was confirmed by flow cytometry for CD3 AF594.

##### T cell culture and activation

Purified CD4+CD25-T cells were cultured in serum-free ImmunoCult™-XF T Cell Expansion Medium (StemCell Tech #10981) supplemented with 30 U/mL rIL-2 (PeproTech 400-02). Cells were activated with Dynabeads® Mouse T-Activator CD3/CD28 at a 1:1 bead-to-cell ratio. Fresh media with IL-2 and beads was changed daily. For co-culture experiments with astrocytes, T cells were washed after 5 days of activation and labeled with CFSE. Labeled T cells were applied directly to washed astrocyte cultures in astrocyte culture medium. Activation and proliferation were assessed via flow cytometry.

#### Cytokine and antigen treatment

Confluent astrocyte cultures were treated with recombinant murine IL1α (3 ng/mL, Sigma-Aldrich, I3901), TNF (30 ng/mL, Cell Signaling Technology, 8902SF), C1q (400 ng/ mL, MyBioSource, MBS143105), and IFNγ (3 ng/mL, abcam ab198568) or IFNβ (1,000 U/mL, Sigma-Aldrich, I8907) for 24 hours to induce a reactive antigen presenting phenotype. For antigen specific OT-II presentation assays, cells were pulsed with endotoxin-free ovalbumin (OVA, 100 µg/mL; InvivoGen) in combination with cytokine cocktail for 24 hours. For antigen specific 2D2 presentation assays, cells were pulsed with endotoxin-free MOG35-55 (10 μg/mL, R&D Systems 2568/1) or whole MOG protein (1 μg/mL, R&D Systems 8536-MO-050) in combination with cytokine cocktail for 24 ho

### Bulk RNA sequencing

RNA was extracted using the RNeasy Plus Mini Kit (Qiagen), and RNA integrity was confirmed by Agilent Bioanalyzer (RIN > 8.4). Libraries were produced using SMART-Seq HT PLUS Kit (Cat. Nos. R400748 &R400749) and sequenced on a NovaSeq X+ at the Genome Technology Center at NYU Grossman School of Medicine at a read depth of 30 million reads per sample. Raw sequencing FastQ data quality was assessed using fastqc^60^. Adapter and quality trimming was performed using trimgalore. Files were then aligned to the mouse genome (GRCm38 - mm10) using salmon (V1.10.0). Differential gene expression testing between conditions was then generated using DESeq2 (V1.42.1)^61^.

### Single cell RNAseq reanalysis

Previously published scRNAseq datasets from mouse cortical astrocytes^4^ were downloaded from GEO ([GSE148611]). Raw or processed expression matrices were reanalyzed using SCTTransform in Seurat (v4) in R. Data were log-normalized, and clustering was performed using principal component analysis (PCA) followed by UMAP for dimensionality reduction. Marker gene expression and module scores for reactive astrocyte signatures were computed using AddModuleScore/.

### Immunocytochemistry

Astrocytes were plated on 96-well glass bottom plates (Revvity 6055300) at 3k/well density and ultimately fixed with 4% paraformaldehyde for 15 minutes at room temperature. Cells were permeabilized with 0.3% Triton X-100 PBS for 45 minutes, blocked with 20% normal donkey serum (Abcam ab7475) in 0.3%Triton X-100 PBS for 45 minutes at room temperature, and incubated overnight at 4 °C with primary antibodies (see Table 1 Key Resources Table) in 0.3% Tri-ton-X. If needed, secondary antibodies were applied for 4 hours at room temperature. Imaging was performed on an inverted Zeiss LSM800 with AiryScan confocal microscope. For in vivo experimentation, mice were euthanized with isoflurane and brains were post-fixed overnight in BD ICC fixation buffer (Fisher BDB550010) (1:1 PBS), cryoprotected in 30% sucrose, and embedded in OCT. Coronal sections (30 µm) were cut on a cryostat and processed for immuno-fluorescence using standard protocols. Mounted sections were washed with PBS, dried at 37 ºC, permeabilized with 0.3% Triton X-100 PBS for 45 minutes at room temperature, blocked with 20% normal donkey serum in 0.3%Triton X-100 PBS for 45 minutes at room temperature, and then stained with antibodies in 0.3%Triton X-100 PBS solution overnight at 4 ºC.

**Table 1.** Key Resource Table.

| **Antibodies** | **Source** | **RRID** | **Identifier** | **Dilution** |
| --- | --- | --- | --- | --- |
| Mouse Anti Mouse GFAP AF647 | Biolegend | AB_2734611 | 837512 | 1:100 |
| Rat Anti GFP AF488 | Biolegend | AB_2563288 | 338008 | 1:100 |
| Alexa Fluor® 488 Rat anti-mouse I-A/I-E | Biolegend | AB_493523 | 107616 | 1:100 |
| Alexa Fluor® 594 Rat anti-mouse I-A/I-E | Biolegend | AB_2566438 | 107650 | 1:100 |
| Alexa Fluor® 647 Rat anti-mouse CD86 | Biolegend | AB_493464 | 105020 | 1:100 |
| Alexa Fluor® 594 Armenian Hamster anti-mouse CD80 | Biolegend | AB_2832338 | 104754 | 1:100 |
| CXCL10 Recombinant Rabbit anti mouse Monoclonal | Thermo | AB_2532429 | 701225 | 1:200 |
| Alexa Fluor® 594 Rat anti-mouse CD3 Antibody | Biolegend | AB_2563427 | 100240 | 1:100 |
| Anti-MHC Class II antibody [6C6] | Abcam | ab55152 | ab55152 | 1:300 |
| Rabbit anti Glial Fibrillary Acidic Protein (Concentrate) | Agilent | Z0334 | Z033401-2 | 1:500 |
| **Other** |  |  |  |  |
| CellTrace™ CFSE Cell Proliferation Kit, for flow cytometry | Thermo |  | C34554 | 1:1000 |
| eBioscience™ Fixable Viability Dye eFluor™ 450 | Thermo |  | 65-0863-18 | 1:1000 |

### Image quantification and colocalization analysis

Fluorescence images of cultured cells were analyzed using QuPath (version 0.7.0). Nuclei were identified by automated detection of DAPI signal using the cell detection algorithm. Cells positive for MHC-II, PDGFRA, or Ki-67 were identified based on marker-specific fluorescence intensity thresholds that were established using negative control samples and applied consistently across all images. Colocalization was determined by the presence of marker-positive signal within the boundaries of DAPI-positive nuclei. The total number of DAPI-positive nuclei and the number of MHC-II+, PDGFRA+, or Ki-67+ cells were quantified for each image. All images were analyzed using identical detection parameters and intensity thresholds across experimental groups.

Fluorescence images of mouse and human tissues were analyzed in Fiji/ImageJ. DAPI staining was used to identify nuclei, while MHC-II- and GFAP-positive signals were thresholded to generate binary masks. Colocalized MHC-II+GFAP+ astrocytes were identified from overlapping thresholded signals and quantified based on associated DAPI-positive nuclei. Total tissue area was determined using a binary tissue mask (MATLAB), and cell counts were normalized to tissue area (cells/mm^2^). For human tissue samples, the shortest distance from each astrocyte to the cortical border was measured in Fiji/ImageJ following manual delineation of the cortex boundary.

### Flow cytometry

T cell-astrocyte cocultures were labeled with efluor 450 viability dye for 40 minutes at 37ºC and then removed from plate with TrypLE. Cells were washed and filtered and fixed with eBioscience™ IC Fixation Buffer (Invitrogen 00-8222-49), then were incubated with Fc block (anti-CD16/32, BioLegend) for 40 minutes at 4 ºC, stained with fluorescently conjugated antibodies diluted in eBioscience™ Flow Cytometry Staining Buffer (ThermoFisher 00-4222-26), and analyzed using a BD Ze5 flow cytometer. Negative and single stained controls were used for compensation and Live/dead discrimination was performed using Fixable Viability Dye eFluor 450. Data were analyzed with FlowJo (BD Biosciences).

### Cytokine profiling via luminex

Supernatants from cytokine-treated astrocytes were collected and analyzed using a multiplex Luminex Proteome Profiler Mouse XL Cytokine Array (R&D #ARY028) per the manufacturer’s protocol. Cytokine concentrations were determined using a Bio-Rad Gel Doc XR imaging system and analyzed with FIJI ImageJ Analysis software. TACM cytokine concentrations were profiled by EVE Technologies Mouse Cytokine/Chemokine 68-Plex Discovery Assay® Array (MD68) in duplicate analysis.

### Human iPSC generation, differentiation, sequencing, and analysis

#### Cells and donors

Peripheral blood mononuclear cells (PBMCs) were obtained from donors with no history of neurological disease at the Johns Hopkins University Multiple Sclerosis Precision Medicine Center of Excellence with the approval of the Johns Hopkins University Institutional Review Board. All participants provided written informed consent allowing for their cells to be repurposed. Blood was collected in sodium heparin coated vacutainers (BD Biosciences 367874) and PBMCs were isolated using SepMate PBMC Isolation Tubes (StemCell Technologies 85460) by mixing 17.5 mL blood with 17.5 mL PBS then adding to SepMate tube with 15 mL Lymphoprep solution (StemCell Technologies 07851) already added under divider. Tubes were centrifuged at 1200 g for 15 minutes, then separated PBMCs were decanted into new 50 mL conical. PBMCs were washed twice with chilled PBS then resuspended in 1 mL of Iscove’s Modified Dulbecco’s Medium (IMDM) supplemented with 10% human serum (Millipore-Sigma H4522-100ML), 100 U/mL penicillin-strepomycin (Thermo Fisher 15140122), 1X GlutaMAX (Thermo Fisher 35050061), and 100 μg/mL Gentamicin (Thermo Fisher 15750060). Cells were then frozen in 10% DMSO at -80 °C in Mr.Frosty (Thermo Fisher 5100-0036) overnight before being transferred to vapor phase liquid nitrogen the next day for long term storage.

#### Automated Reprogramming of PBMCs to iPSCs

PBMCs were thawed and recovered overnight in Stem-Pro-34 SFM Complete Medium (Thermo Fisher, 10639-011) supplemented with SCF (R&D Systems, 255-SC 200 ng/mL), Flt3 (Thermo Fisher Scientific, 35050-079 200 ng/mL), IL3 ( R&D Systems, 203-IL 40 ng/mL), IL6 (R&D Systems, 206-L 40 ng/mL), and GlutaMAX (Thermo Fisher 35050-061). The following day, PBMCs were seeded at 60K and 100K densities onto 96-well flat bottom plates coated with Cultrex (HESC qualified, R&D Systems 3434-010-02) for reprogramming. Reprogramming was performed using the CytoTuneiPS Sendai Reprogramming v2.0 Kit (Thermo Fisher A16517) per the manufacturer’s recommendations, with modifications for cell number and plate format. Post-infection, cells were gradually transitioned to Freedom media (DMEM-F12 with Freedom-1 Supplement, Life Technologies, Custom) for 5 days, with daily media changes on the NYSCF Global Stem Cell Array platform^62,63.^ Tra-1-60 live cell surface staining was used 12–14 days post-transfection to identify reprogrammed cells. Successfully reprogrammed lines were consolidated in a Cultrex-coated 96-well plate and cryostored in LN2 upon reaching confluency. Sendai reprogrammed iPSCs were enriched by FACS sorting (CCD56-, CCD13-, Tra-1-60+, SSEA4+ cells) and then monoclonalized^64^. Monoclonalized iPSC lines were expanded on the automated NYSCF Global Stem Cell Array® platform and then frozen in Synth-a-Freeze Cryopreservation Media (ThermoFisher Scientific, A1254201) at 500K cells/vial. All iPSC lines underwent a stringent quality control, encompassing sterility and mycoplasma checks, viability, karyotyping (Illumina Global Screening Array), SNP ID fingerprinting (Fluidigm SNPTrace), pluripotency and embryoid body scorecard assays (NanoString). For differentiations toward CNC cells, iPSCs were thawed onto 6-well Cultrex-coated plates and adapted to mTeSR1 medium (StemCell Technologies 85850). For passaging cells were dissociated via enzymatic digestion with StemPro Accutase (Thermo Fisher Scientific 00-4555-56) for 3-5 minutes and re-plated in mTeSR1 medium with the addition of 10µM Y-27632 (AbMole M1817) for 24 hours.

#### iPSC differentiation into glial-enriched cultures (astrocytes and oligodendrocytes)

Glia-enriched cultures, containing both astrocytes and oligodendrocytes, were generated using a previously established protocol^53,65^. Briefly, iPSCs were seeded on Geltrex-coated plates and cultured in Neural Induction medium for eight days, followed by N2 medium for four days. On day 12, confluent wells were scraped, and resulting clumps were gently triturated and transferred into ultra-low-attachment (ULA) 6-well plates. These were then fed every other day with N2B27 Medium until day 20, at which point the medium was switched to PDGF medium. At day 30 spheres with a dark golden core, measuring from 300–600 µm, were selected, seeded into poly-L-ornithine/laminin (Sigma Aldrich P3655; Gibco 23017-015) coated plates and fed every other day with PDGF medium. From day 30 progenitors migrated out of the spheres and differentiated into neurons, astrocytes and oligodendrocyte lineage cells. Cultures were kept until day 70 when they were dissociated for single cell RNA sequencing analysis.

The composition of each sequential medium used is follows: Neural Induction medium (d0-d7): mTeSR1 without selected factors (StemCell Technologies 05892), Pen-Strep (Gibco 15070063 1:100), SB431542 (Selleck Chemicals, S1067 10 µM) LDN193189 (ReproCELL 04-0074 250 nM), all trans retinoic acid (RA, Sigma-Aldrich R2625 100 nM). N2 Medium (d8-d11): DMEM/F12 + GlutaMAX (Gibco 10565042), N2 supplement (Gibco 17502048 1:100), RA, smoothened agonist (EMD Millipore 566660 1 µM), 2-Mercaptoethanol (Thermo Fisher Scientific 21985023 1:1000), MEM non-essential amino acids (Gibco, 11140050 1:100). N2B27 Medium (d12-d19): N2 medium, B27 supplement without vitamin A (Thermo Fisher Scientific 12587010 1:50), insulin solution (Millipore Sigma, I9278-100ml 25 µg/mL). PDGF Medium (d20-d70):DMEM/F12 + GlutaMAX, Pen-Strep, 2-Mercaptoethanol, MEM non-essential amino acids, N2 supplement, B27 supplement without vitamin A, insulin solution, PDGFaa (R&D Systems, 221-AA 10 ng/mL), IGF-1 (R&D Systems, 291-G1 10 ng/mL), HGF (R&D Systems, 294-HG 5 ng/mL), NT3 (R&D Systems, 267-N3-MTO 10 ng/ mL), biotin (Sigma-Aldrich, B4639 100 ng/mL), N6,2-O-dibutyryladenosine 3,5 -cyclic monophosphate sodium salt (db-CaMP, Sigma Aldrich, D0627 1 µM).

#### iPSC differentiation into microglial progenitor cells and subsequent incorporation into glial-enriched cultures

Microglia progenitor cells were generated by adapting a previously established protocol^66^. For mesodermal induction iPSCs were passaged once in mTeSR™1 medium with Revi-taCell supplement (Thermo Fisher Scientific A264450) then plated into a 96-well ultra-low attachment U-bottom plate (1*104 cells/well) for embryoid body (EB)-like aggregation in mTeSR™1 medium supplemented with BMP4 (R&D Systems, 314-BP-MTO 50 ng/mL), VEGF (R&D Systems 293-VE 50 ng/mL) and SCF (R&D Systems, 255-SC 20 ng/mL). On day 4 EBs were transferred to 6-well plates and fed with X-VIVO 15 medium supplemented with SCF (R&D Systems 255-SC, 50 ng/mL), M-CSF (R&D Systems, 216-MC 50 ng/ mL), IL-3 (R&D Systems, 203-IL 50 ng/mL), Flt3-ligand (Thermo Fisher Scientific 35050-079 50 ng/mL) and TPO (R&D Systems, 288-TP 5 ng/mL). For final differentiation to microglial progenitors, on day 11 medium was switched to X-VIVO 15 supplemented with Flt3-ligand (50 ng/mL), M-CSF (R&D Systems 216-MC 50 ng/mL) and GM-CSF (R&D Systems 215-GM 25 ng/mL). Between day 11 and day 18, released microglial progenitors accumulated in the supernatant. On day 18, supernatants were filtered into a 50 ml conical tube and isolated microglial progenitors were added to day 50 glial-enriched cultures. Cultures were maintained in PDGF medium, with the addition IL-34 (R&D Systems 5625-IL 100 ng/ mL) and GM-CSF (R&D Systems 215-GM 10 ng/mL) to support microglial maturation. Following microglial integration, a two-thirds media change was performed carefully from the side to avoid lifting or disrupting the cultures.

#### Stimulation of iPSC-derived mixed cultures with inflammatory cytokines and dissociation to single cell suspension

On day 68 glial-enriched cultures with incorporated microglia were exposed to TNF (Peprotech AF-300-01A 50 ng/ mL) and IFNy (Peprotech AF-300-02 20 ng/mL) for 48h before proceeding to single cell dissociation. Day 70 cultures were dissociated with papain (Worthington Biochemical, LK003150), filtered into 50 mL conical tubes using 40 µm strainers as previously described^53^, and counted for single cell RNA sequencing.

#### iPSC-derived mixed culture single cell transcriptomics library preparation and sequencing

Single cell transcriptomic analysis of iPSC-derived mixed cultures was performed using 10x Genomics Chromium Next GEM Fixed RNA Profiling (also known as 10x Flex) Reagent Kits according to manufacturer’s recommendations^67^. At day 70 in culture, 48 hours after the addition of cytokine, single cells were collected from wells via papain digestion, washed, and then fixed and permeabilized overnight at 4 °C in 1 mL fixation buffer (10% Concentrated Fix and Perm Buffer (10x Genomics PN-2000517), 4% formaldehyde (Fisher Bioreagents BP351-25) in nuclease free water). The next day, cells were centrifuged then resuspended in 1 mL quenching buffer (12.5% Concentrated Quench Buffer, 10x Genomics PN-2000516 in nuclease free water) followed by the addition of 0.1mL Enhancer (10x Genomics PN-2000482) and stored for 2 days at 4 °C. Glycerol was then added to a concentration of 10% and samples were frozen at -80 °C until both batches were completed. Samples were thawed together and washed in 1mL Phosphate Buffered Saline (PBS) with 0.02% Bovine Serum Albumin (BSA). Cells were counted on a Countess 3 automated cytometer with Acridine Orange/Propidium Iodide then pelleted and underwent barcoded probe hybridization using Chromium Fixed RNA Profiling reagents as specified by manufacturer (10x Genomics CG000527, Rev F). Briefly, up to 2×106 fixed and permeabilized cells were incubated at 42 °C for approximately 19 hours in the presence of barcoded probes targeting 18,129 genes. Following hybridization, samples were again counted and then equal numbers of cells per sample (one sample = one well from one donor treated with one condition) were combined into 4 distinct pools. For each donor (N = 6), all conditions for that donor were in the same pool (N = 3: Vehicle, IFNγ, TNF) except the one technical replicate in which a single TNF treated sample was split into two for probe hybridization. The technical replicate was run in a different pool than the other 3 samples from that donor. Pooled cells were then washed three times, counted, and then loaded onto 10x Genomics microfluidics Chip Q (PN-1000418) targeting 8,000 cells per sample and partitioned into individual gel bead-in-emulsion (GEMs) on a Chromium iX. Following partitioning, GEMs were incubated in a thermal cycler to allow for probe ligation and addition of UMI and GEM barcode. GEMs were then broken followed by pre-amplification and addition of Illumina sequencing adapters (1 per pool, using a total of 9 PCR cycles). The 4 libraries were then sequenced on an Illumina NovaSeq X Plus in a 28×10×10×90 configuration at the Genetics Resources Core Facility RRID SCR_018669, Johns Hopkins Department of Genetic Medicine, Baltimore, MD.

#### iPSC-derived mixed culture single cell transcriptomics data analysis

Transcript data was demultiplexed into individual samples and quantified with Cellranger Multi (10x Genomics, v9.0.1) using Human Transcriptome Probe Set v1.1.0 and the GRCh38-2024-A reference genome expecting 8,000 cells per sample. Single cell analysis was then performed according to previously published best practices^68,69^. Filtered barcode-feature matrices (155,392 cells across 19 samples) from Cell-ranger were imported into R (v4.5.1). Prior to any filtration, likely doublets were identified using scDblFinder (v1.22.0)^70^. Predicted doublets were retained through later processing stages to help identify doublet-enriching clusters. An initial filtering step was then applied to remove the most obvious low-quality cells which had UMI counts less than 500, feature counts (total detected genes) less than 200, or percent of reads mapping to mitochondrial transcripts greater than 5%. This filtering step left 154,971 cells. Library sizes were then normalized with Seurat (v5.3.0) NormalizeData function^71^. Variable features per condition were identified with Seurat’s VariableFeatures function and the union of these features were then scaled with ScaleData followed by principal component analysis. To facilitate identifying shared cell types across conditions, in the first round of analysis the three conditions were integrated with Harmony^72^. Non-linear dimensionality reduction was performed with uniform manifold approximation and projection (UMAP) followed by k-nearest neighbor graph computation and Louvain unsupervised clustering. Using marker gene expression, clusters were labeled as being primarily astrocytes, oligodendrocyte precursor cells (OPCs), oligodendrocytes, neurons, or microglia. In addition to these primary cell types, two additional clusters were identified and removed from downstream analysis. The first was characterized by gene expression consistent with cellular response to hypoxia. The second expressed genes associated with mesenchymal lineage cells^73^. Both contaminating cell types were present in approximately equal proportions across all three conditions. For all stages in the analytic pipeline other than this initial stage, integration was performed on the donor level rather than the condition so that we could focus on the differences caused by cytokine treatment rather than between-person variation.

Cells were then analyzed in a hierarchical fashion wherein the clusters identified as belonging to each cell type were then subsetted and each cell type then underwent a second round of quality control where outlier cells were identified as either having mitochondrial percentages 3 median absolute deviations greater than the sample’s cell-type median, or library sizes (UMI counts or feature counts) 3 median absolute deviations less than the sample’s cell-type median (on a log_2_ scale). These calculations were performed with the isOutlier function from the scuttle package^74^. Outlier detection was run at this stage in a cell-type specific fashion rather than in the initial stage with mixed cell types to avoid either outlier under or over-detection caused by differences in cell metabolism or library size between different cell types (i.e. astrocytes vs. neurons). Following outlier detection, the above analysis pipeline was repeated on each cell type: library size normalization, variable feature detection (run for each donor and then combined) and scaling, principal component analysis, integration with Harmony (integrating on donor), k-nearest neighbor graph calculation, non-linear dimensionality reduction and unsupervised clustering. Within each cell type, clusters were discarded when they appeared to be doublets (enriching for cells predicted to be doublets by scDblFinder and/or expressing high levels of genes associated with other cell types), were characterized by either very low library size or high mitochondrial percentages or were characterized by cellular response to hypoxia. All three kinds of poor-quality clusters were present in equal proportions across condition and donor. Following the removal of poor-quality clusters for each major cell type, cleaned cells from each cell type were recombined (total of 110,342 remaining cells) and reprocessed, again integrating on donor, followed by non-linear dimensionality reduction with UMAP as shown in Figure 5B. Each major expected cell type (astrocytes, OPCs, oligodendrocytes, neurons, microglia) were present. In addition to the primary cell types, we identified proliferative subtypes in the astrocyte, OPC, and microglia populations (identified as proliferative on Figure 4B, C). We also identified a small subset of astrocyte-like cells characterized by expression of genes associated with cilium assembly and function (labeled Astrocytes – Cilia on Figure 4B, C). This population was present across all 3 conditions and was distinct from the other astrocytes populations (while still retaining high expression of astrocyte identity genes such as GFAP, AQP4, and SPARCL1 and having no significant expression of markers associated with other cell types). Throughout the analysis process we utilized the scCustomize package (v3.2.0) for QC metric calculation and data visualization^75^.

To test for differential gene expression between conditions (Vehicle, IFNγ, TNF) we performed pseudobulking analysis with the non-proliferative astrocytes (not including the sub-type characterized by cilia formation). For each independent sample, raw counts for all cells belonging to the cell type were aggregated into a single pseudobulk sample. There were no instances of pseudobulk samples being created from less than 10 cells. The single technical replicate was not used in the pseudobulk analysis. We then performed bulk RNA seq analysis using the pseudoBulkDGE function from the scran package^76^ which is a wrapper function around edgeR’s quasi-likelihood framework for differential expression testing^77^. We included donor as a blocking factor in the generalized linear model (design = ∼ 0 + donor + condition) to control for between person variability and instead focus on changes induced by the different cytokine treatments. We tested three pairwise contrasts: TNF vs. Vehicle, IFNγ vs. Vehicle, and IFNγ vs. TNF. For the purposes of gene ontology enrichment testing, genes were considered to be differentially expressed if they had a greater than 1.5-fold change in either direction and an adjusted p-value less than 0.05. We then performed hypergeometric analysis on the upregulated and downregulated genes with the enrichGO function from the clusterProfiler package (v4.16.0)^78^. Volcano plots from pseudobulk analyses in Figure 5 were generated with the package EnhancedVolcano (v1.26.0)^79^.

## ACKNOWLEDGEMENTS

The computational requirements for this work were supported in part by the NYU Langone High Performance Computing (HPC) Core’s resources and personnel. Funding for this work was provided by NIH/ NEI (R01EY033353), the Cure Alzheimer’s Fund, the MD Anderson Belfer Neurodegeneration Consortium, Howard Hughes Medical Research Institute, Howard Hughes Medical Institute Emerging Pathogens Initiative (S.A.L., A.I.), and the Carol and Gene Ludwig Family Foundation (S.A.L.). The iPSC-based work was supported by the Bloomberg Philanthropies via the JHU-NYSCF Precision Medicine Partnership (PI Antony Rosen, Dean of Research, JHU). We would also like to thank the Leon Levy Foundation’s Fellowship in Neuroscience for their support (F.L.). We thank the NYSCF Global Stem Cell Array® Team for generating the iPSC lines. The iPSC-based work was supported by the Bloomberg Philanthropies via the JHU-NYSCF Precision Medicine Partnership (PI Antony Rosen, Dean of Research, JHU).

## AUTHOR CONTRIBUTIONS

T.M.F. and S.A.L. conceived the study and wrote the manuscript. All authors reviewed and agreed to the final manuscript. T.M.F., F.L., and C.W. performed cell culture and bulk RNA sequencing and analysis. T.M.F. and M.D.S. performed data analysis on published scRNAseq. T.M.F. and K.L. planned and performed immunofluorescence on brain sections and astrocyte/T cell cultures. P.K. set up, optimized, and collaborated on mouse EAE experiments. N.P. and J.M performed human iPSC differentiations to mixed CNS cultures. M.D.S., V.F., and P.A.C. planned, executed, and analyzed human iPSC experiments. S.A.L., A.I., and P.A.C. obtained funding.

---

**Supplemental Figure S1.**
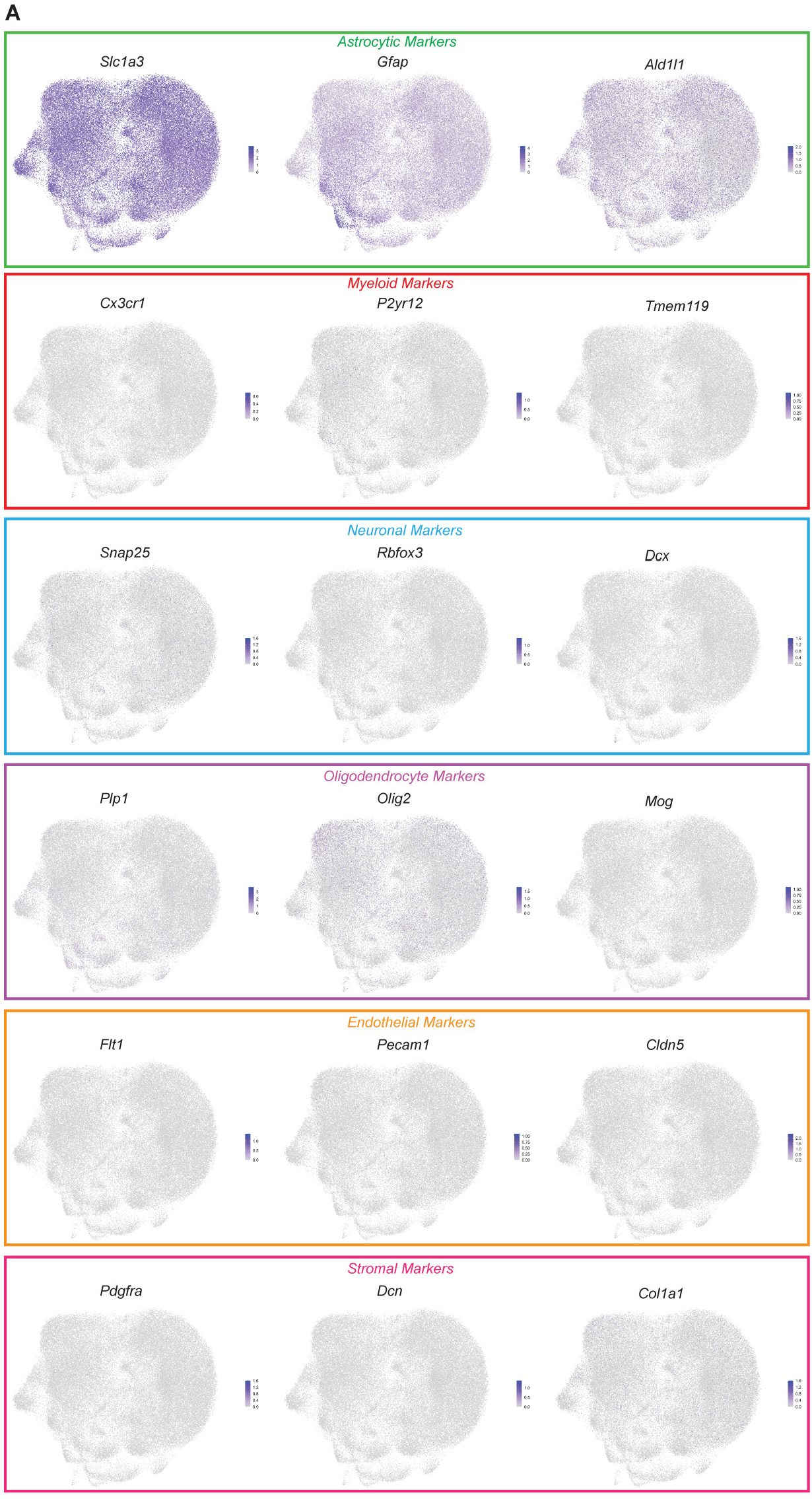
Clustered cell populations do not express other cell type transcripts. **A**. Expression of genes associated with myeloid cells (e.g. *Cx3cr1, P2yr12, Tmem112*), neurons (e.g. *Snap25, Rbfox3, Dcx*), oligodendrocyte lineage cells (e.g. *Plp1, Olig2, Mog*), endothelial cells (e.g. *Flt1, Pecam1, Cldn5*), or stromal cells (e.g. *Pdgfra, Dcn, Col1a1*) was not measurable in any cell identified as ‘astrocytes’.

**Supplemental Figure S2.**
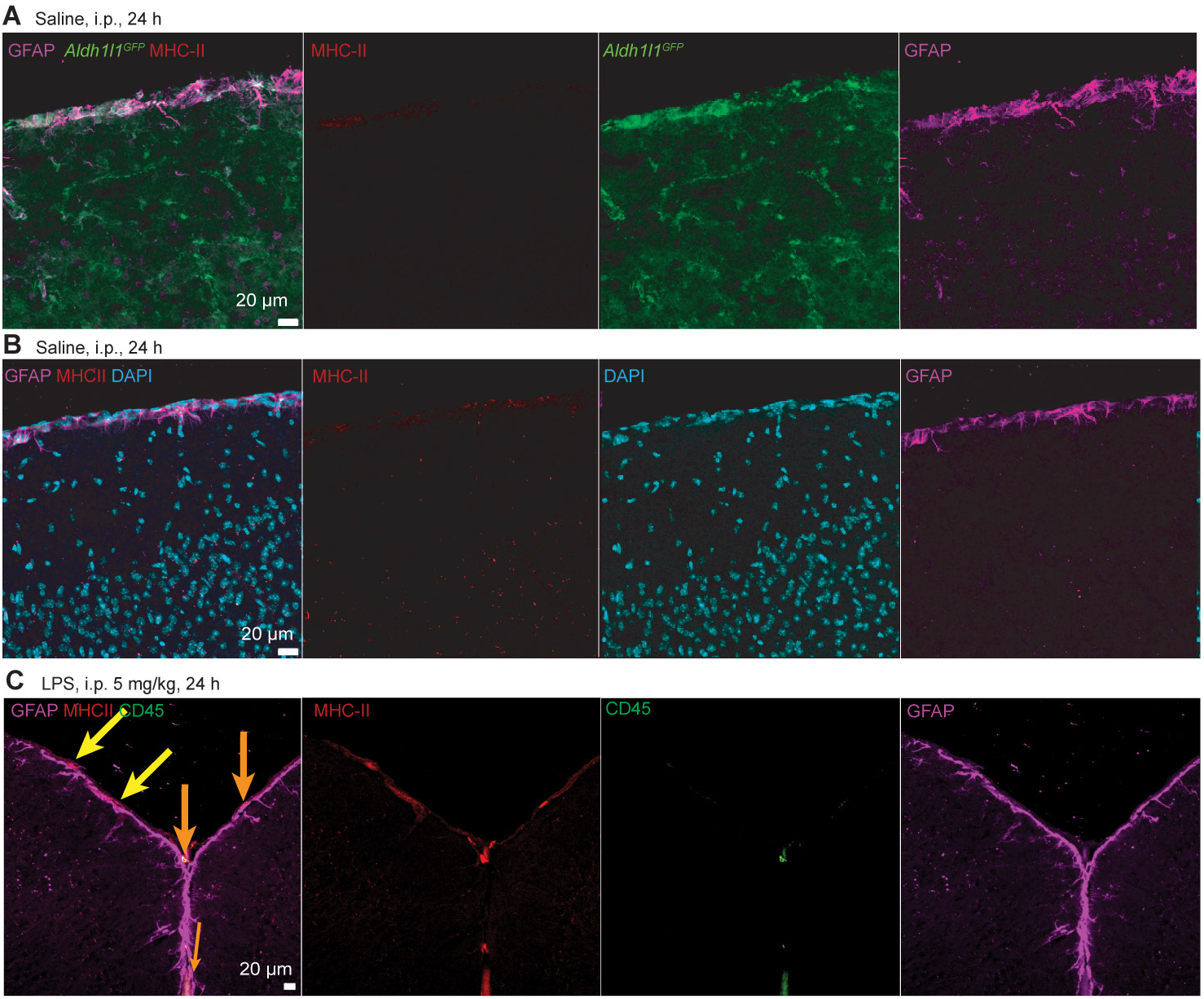
MHC-II+GFAP+CD45-cells do not appear at the brain surface in saline injected controls. **A**. Representative micrograph of surface astrocytes (GFAP, magenta; *Aldh1l1*^eGFP^, green) co-labelling for MHC-II (red) after 24 hours of Saline injection i.p. in *Aldh1l1*^eGFP^+ mice. **B**. Representative micrograph of surface astrocytes (GFAP, magenta) co-labelling for MHC-II (red) after 24 hours of Saline injection i.p. **C**. Representative micrograph distinguishing between MHC-II+GFAP+CD45-astrocytes (yellow) and MHC-II+CD45+ immune cells (orange) at the cortex border 24 hours after i.p. LPS injection.

**Supplemental Figure S3.**
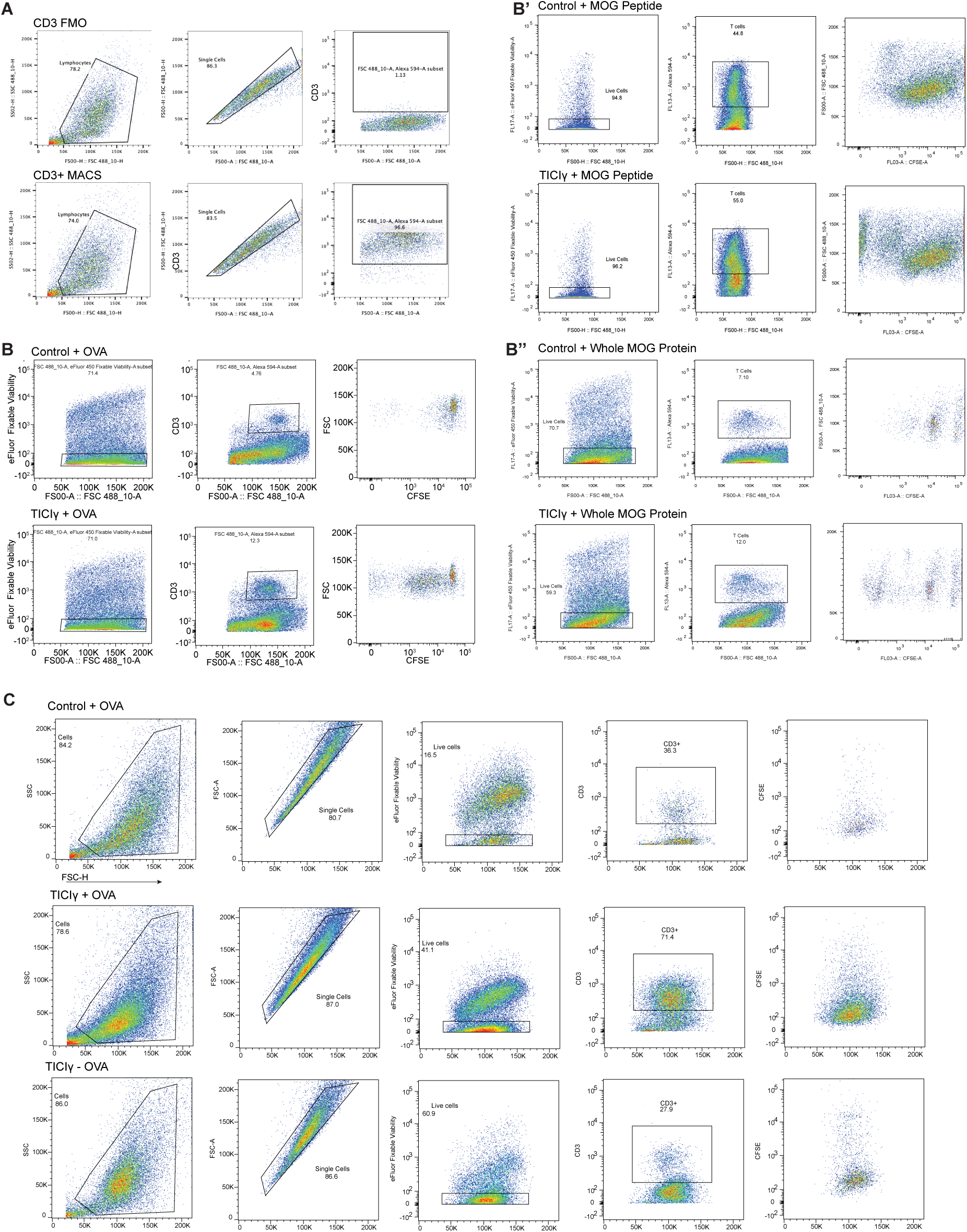
Flow Cytometry gating and cytokine measurements in T cell-Astrocyte coculture. **A**. Flow cytometry gating strategy for MACS-purified T cells. Single cells are gated based on forward and side scatter, subsequently with CD3-AlexaFluor 594. Negative staining for CD3 (top) shows no contribution to T cell gating numbers while positive staining (bottom) shows high T cell purity after MACS purification. **B-B’’**. Representative flow cytometry gating for CD3+ OT-II or 2D2 T cells and astrocytes after coculture. Expansion of both CD3 positive T cells and negative astrocytes are seen after TICIγ pretreatment (bottom) as compared to when no cytokines are applied (top). **C**. Representative flow cytometry gating for CD3+ T cells after naïve T cell astrocyte coculture. Conditions shown are with cytokine and OVA (top), with cytokine no OVA (middle) and no cytokines with OVA (bottom). X axis for all plots is FSC-H.

**Supplemental Figure S4.**
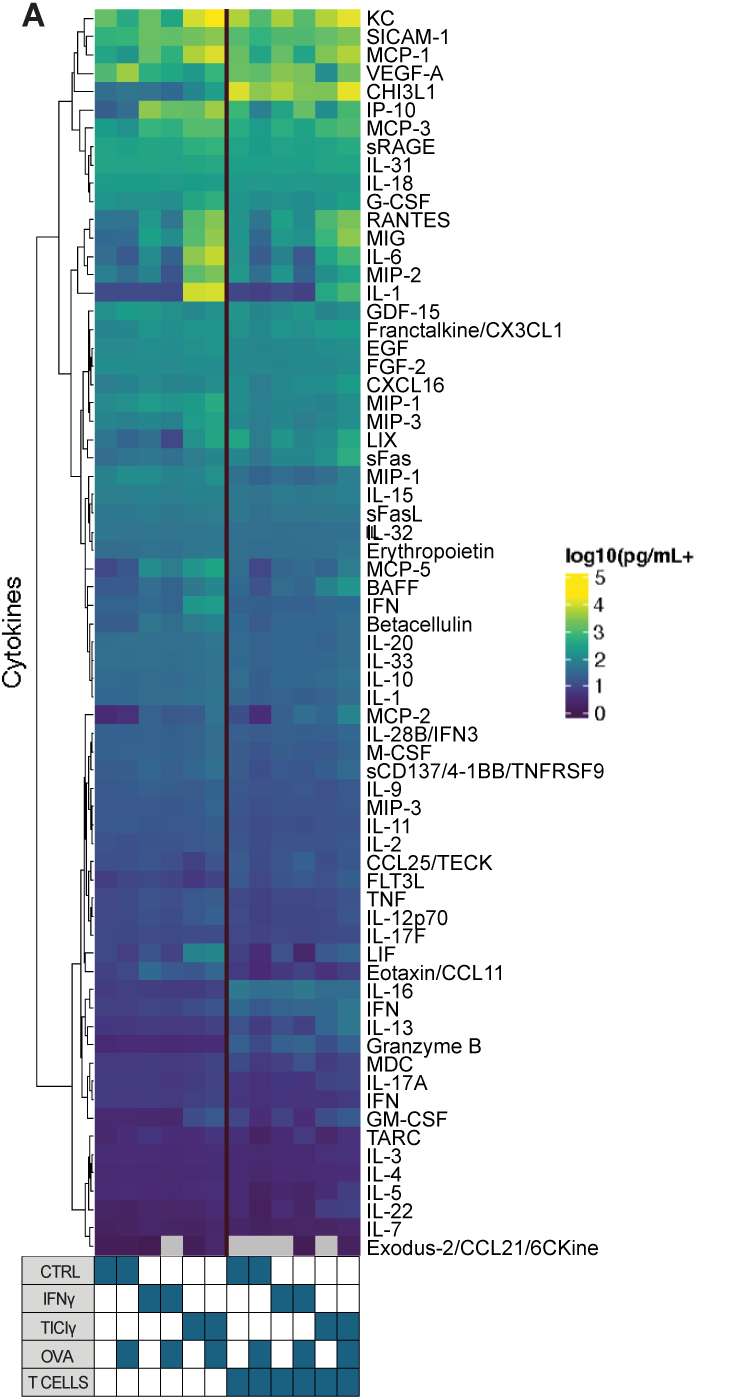
Heatmap of cytokine concentrations in ACM and TACM.

**Supplemental Figure S5.**
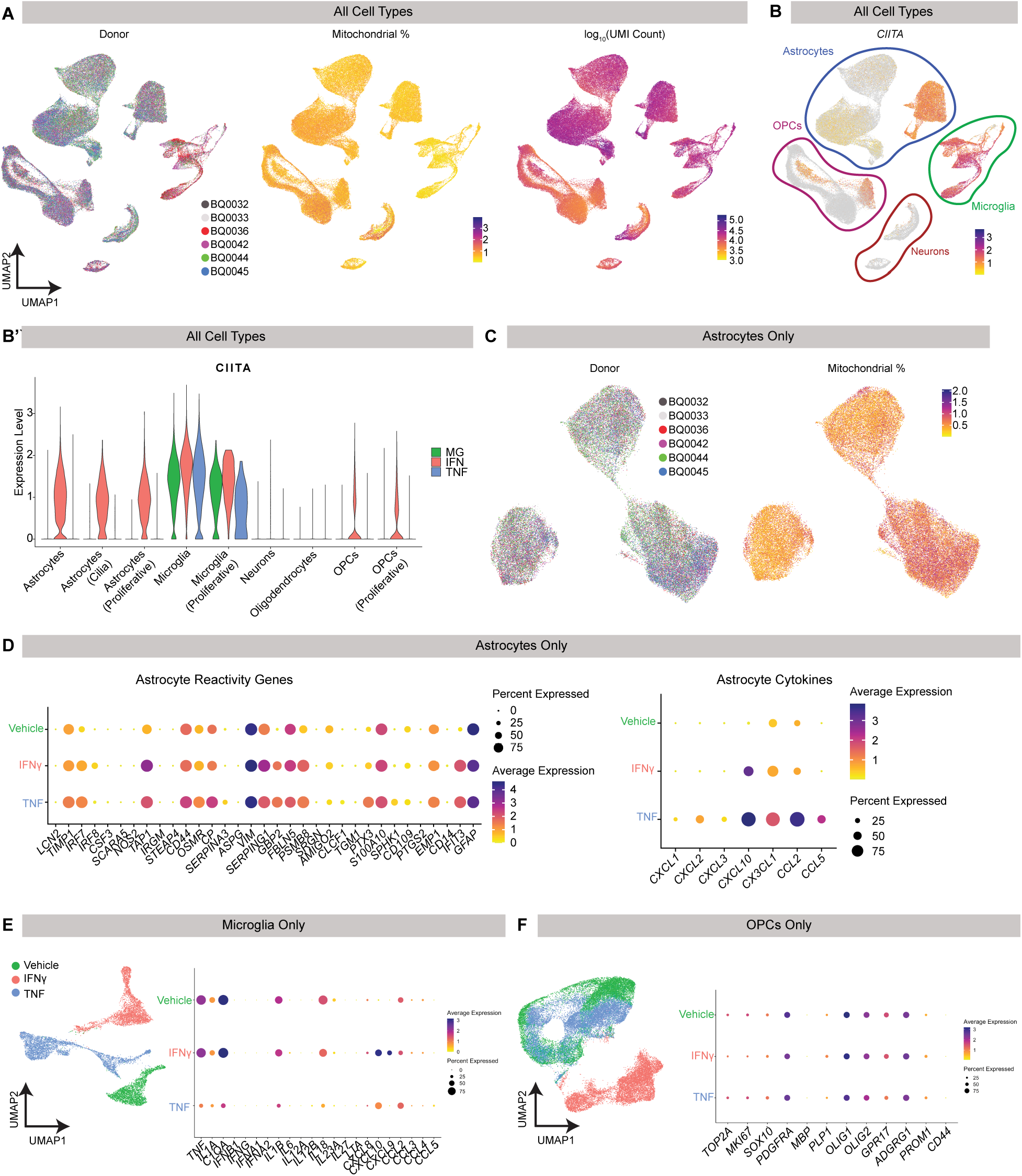
Extended analysis of human iPSC data. **A**. UMAP plotting of all sequenced cells split by patient ID and feature plotting of mitochondrial gene expression % and UMI contribution. N = 6 individual patient iPSC lines were differentiated, cultured, treated, and sequenced. **B**. Feature and violin plotting of CIITA gene expression across all cell types. **C**. UMAP plotting of subsetted astrocytes split by patient ID and feature plotting of mitochondrial gene expression %. **D**. Dot plotting of subsetted astrocyte gene expression across conditions of reactivity genes (left) and relevant cytokine genes (right). **E**. UMAP and dot plotting of subsetted microglia across conditions. Genes included are specific to known microglial-derived inflammatory cytokines. **F**. UMAP and dot plotting of subsetted OPCs across conditions. Genes included are specific to proliferation and lineage commitment.

**Supplemental Figure S6.**
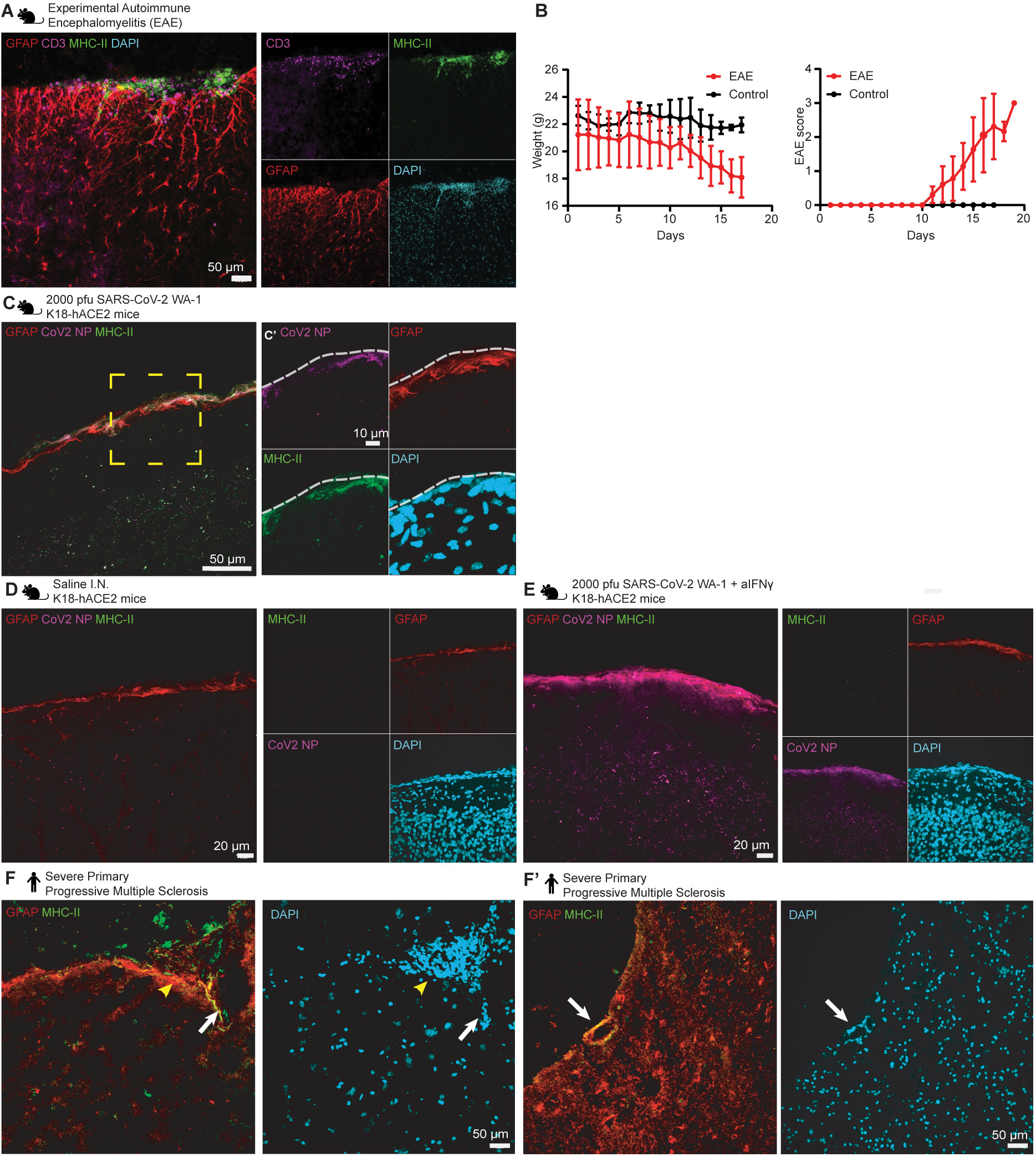
In vivo examples of surface astrocytes expressing MHC-II. **A**. Imaging of lumbar spinal cord lesion in EAE induced mouse at peak EAE score 3. Astrocytes (GFAP, red) are in close proximity to hypercellular lesion showing increased MHC-II cells (green) and CD3+ T cells (magenta). **B**. Plotting of EAE scores and body weight changes over time course of EAE. (bars = SEM) (N = 10 EAE, 5 CTRL). **C**. Representative micrograph of surface astrocytes (GFAP, red) co-labelling for MHC-II (green), infected with SARS-CoV2 nucleocapsid (magenta). Image collected 5 days after SARS-CoV2 inoculation in mice intranasally (top) or intranasal saline administration (bottom). **C’**. (higher magnification of box in C). **D-E**. Representative micrograph of surface astrocytes (GFAP, red), labelling for MHC-II (green), SARS-CoV2 and nucleocapsid (magenta). Image collected 5 days after saline or (D) SARS-CoV2 + -IFNγ blocking antibody i.p. (E). **F**. Representative micrograph of human astrocytes (GFAP, red) and labelling for MHC-II (green) near large vasculature (F’) and hypercellular pathology. Images collected in human pre-frontal during severe primary progressive multiple sclerosis (see Table S2 for patient details).

**Supplemental Table S1.** Donor demographics for human iPSC cultures.

| Pool | Donor | Sex | Race | Batch |
| --- | --- | --- | --- | --- |
| A | BQ0044 | F | W | 1 |
| A | BQ0033 | F | B | 1 |
| B | BQ0036 | F | B | 2 |
| B | BQ0032 | F | B | 2 |
| C | BQ0045 | M | W | 2 |
| D | BQ0042 | F | W | 1 |

**Supplemental Table S2.** Patient sample details for human postmortem immunostaining.

| Multiple Sclerosis human tissue |  |  |  |  |
| --- | --- | --- | --- | --- |
| Sample ID | Age Year | Sex | Lesion Type | Pathology |
| 5909 | 62 | Woman | Type III | MS |
| 5949 | 75 | Man | Type III | MS |
| 5997 | 75 | Man | Type III | MS |
| 6097 | 53 | Man | Type III | MS |
| 6127 | 71 | Man |  | Non-MS |
| 6148 | 62 | Woman |  | Non-MS |
| 6302 | 66 | Woman |  | Non-MS |
| 6300 | 68 | Man |  | Non-MS |
| Alzheimer's Disease Human tissue |  |  |  |  |
| DONOR_NUMBER | SAMPLE_ID | AGE | SEX | PATHOLOGY |
| D18(IF) | TN11-23 | 74 | M | Alzheimer's disease-severe |
| D19(IF) | TN11-45 | 69 | M | Alzheimer's disease-severe |
| D20(IF) | TN12-49 | 80 | F | Alzheimer's disease-severe |
| D21(IF) | TN12-60 | 92 | M | Alzheimer's disease-severe |
| D22(IF) | TN16-37 | 92 | F | Alzheimer's disease-severe |
| D23(IF) | TN17-49 | 94 | M | Alzheimer's disease-severe |
| D24(IF) | TN13-66 | 95 | F | Non-Alzheimer's disease |
| D26(IF) | TN13-11 | 88 | M | Non-Alzheimer's disease |
| D27(IF) | TN13-21 | 49 | F | Non-Alzheimer's disease |
| D28(IF) | TN14-74 | 46 | M | Non-Alzheimer's disease |
| D289(IF) | TN18-34 | 61 | F | Non-Alzheimer's disease |

